# Variably methylated regions in the newborn epigenome: environmental, genetic and combined influences

**DOI:** 10.1101/436113

**Authors:** Darina Czamara, Gökçen Eraslan, Jari Lahti, Christian M. Page, Marius Lahti-Pulkkinen, Esa Hämäläinen, Eero Kajantie, Hannele Laivuori, Pia M Villa, Rebecca M. Reynolds, Wenche Nystad, Siri E Håberg, Stephanie J London, Kieran J O’Donnell, Elika Garg, Michael J Meaney, Sonja Entringer, Pathik D Wadhwa, Claudia Buss, Meaghan J Jones, David TS Lin, Julie L MacIsaac, Michael S Kobor, Nastassja Koen, Heather J Zar, Karestan C Koenen, Shareefa Dalvie, Dan J Stein, Ivan Kondofersky, Nikola S Müller, Fabian J Theis, Major Depressive Disorder Working Group of the Psychiatric Genomics Consortium, Katri Räikkönen, Elisabeth B Binder*

**Author notes:** A detailed author list is given in the Supplemental Text. <u>Correspondence:</u> Elisabeth Binder, MD PhD, Department Translational Research in Psychiatry, Max Planck Institute of Psychiatry, Kraepelinstr. 2, 80804 Munich, Germany.

## Abstract

**Background:** Epigenetic processes, including DNA methylation (DNAm), are among the mechanisms allowing integration of genetic and environmental factors to shape cellular function. While many studies have investigated either environmental or genetic contributions to DNAm, few have assessed their integrated effects. We examined the relative contributions of prenatal environmental factors and genotype on DNA methylation in neonatal blood at variably methylated regions (VMRs), defined as consecutive CpGs showing the highest variability of DNAm in 4 independent cohorts (PREDO, DCHS, UCI, MoBa, N=2,934).

**Results:** We used Akaike’s information criterion to test which factors best explained variability of methylation in the cohort-specific VMRs: several prenatal environmental factors (E) including maternal demographic, psychosocial and metabolism related phenotypes, genotypes in cis (G), or their additive (G+E) or interaction (GxE) effects. G+E and GxE models consistently best explained variability in DNAm of VMRs across the cohorts, with G explaining the remaining sites best. VMRs best explained by G, GxE or G+E, as well as their associated functional genetic variants (predicted using deep learning algorithms), were located in distinct genomic regions, with different enrichments for transcription and enhancer marks. Genetic variants of not only G and G+E models, but also of variants in GxE models were significantly enriched in genome wide association studies (GWAS) for complex disorders.

**Conclusion:** Genetic and environmental factors in combination best explain DNAm at VMRs. The CpGs best explained by G, G+E or GxE are functionally distinct. The enrichment of GxE variants in GWAS for complex disorders supports their importance for disease risk.

## Introduction

Fetal or prenatal programming describes the process by which environmental events during pregnancy influence the development of the embryo with on-going implications for future health and disease. Several studies have shown that the *in utero* environment is associated with disease risk, including coronary heart disease ^1; 2^, type 2 diabetes ^3^, childhood obesity ^4; 5^ as well as psychiatric problems ^6^ and disorders ^7-9^.

Environmental effects on the epigenome, for example via DNA methylation, could lead to sustained changes in gene transcription and thus provide a molecular mechanism for the enduring influences of the early environment on later health ^10^. Smoking during pregnancy influences widespread and highly reproducible, albeit modest in magnitude, differences in DNA methylation at birth ^11^. Less dramatic effects have been reported for maternal folate status ^12^, maternal body mass index (BMI) ^13^, pre-eclampsia, gestational diabetes ^14; 15^, toxins ^16^ and also assisted reproductive technology ^17^. Possible epigenetic changes as a consequence of prenatal stress are less well established ^18^. Some of these early differences in DNA methylation persist, although attenuated, through childhood ^11; 19^ and might be related to later symptoms and indicators of disease risk, including behavioral abnormalities ^14^, impaired lung function ^19^ and BMI during childhood ^20; 21^ or substance use in adolescence ^22^. These data emphasize the potential importance of the prenatal environment for the establishment of inter-individual variation in the methylome as a predictor or even mediator of disease risk trajectories.

In addition to the environment, the genome plays an important role in the regulation of DNA methylation. To this end, the impact of genetic variation, especially of single nucleotide polymorphisms (SNPs) on DNA methylation in different tissues, has resulted in the discovery of a large number of methylation quantitative trait loci (meQTLs, i.e., SNPs significantly associated with DNA methylation status ^23^). These variants are primarily in cis, i.e., at most 1 million base pairs away from the DNA methylation site ^23-25^ and often co-occur with expression QTLs or other regulatory QTLs ^26-28^. The association of meQTLs with DNA methylation is relatively stable throughout the life course ^24^ and meQTLs show some cross-tissue correlation ^27^. In addition, SNPs within meQTLs are strongly enriched for genetic variants associated with common disease in large genome-wide association studies (GWAS) such as BMI, inflammatory bowel disease, type 2 diabetes or major depressive disorder ^24; 26; 27; 29^.

Environmental and genetic factors may act in an additive or multiplicative manner to shape the epigenome to modulate phenotype presentation and disease risk ^30^. However, very few studies have so far investigated the joint effects of environment and genotype on DNA methylation, especially in a genome-wide context. Klengel et al. ^31^ reported an interaction of the FK506 binding protein 5 gene (*FKBP5*) SNP genotype and childhood trauma on *FKBP5* methylation levels in peripheral blood cells, with trauma associated changes only observed in carriers of the rare allele. Chen et al. ^32^ found that a variant in the brain derived neurotrophic factor gene (*BDNF*) strongly moderated the association between antenatal maternal anxiety and DNA methylation at birth. Gonseth et al. ^33^ observed that three of the ten most significant maternal-smoking sensitive CpG-sites in newborns identified in large epigenome wide studies were also significantly associated with SNPs located proximal to each gene. The most comprehensive study of integrated genetic and environmental contributions to DNA methylation so far was performed by Teh et al. ^34^. This study examined variably methylated regions (VMRs), defined as regions of consecutive CpG-sites showing the highest variability across all methylation sites assessed on the Illumina Infinium HumanMethylation450 BeadChip array. In a study of 237 neonate methylomes derived from umbilical cord tissue, the authors explored the proportions of the influence of genotype vs. prenatal environmental factors such as maternal BMI, maternal glucose tolerance and maternal smoking on DNA methylation at VMRs. They found that 75% of the VMRs were best explained by the interaction between genotype and environmental factors (GxE) whereas 25% were best explained by SNP genotype and none by environmental factors alone. After correction for ethnicity, which inflated the influence of genotype, 85% of VMRs were best explained by GxE. Collectively, these studies highlight the importance of investigating the combination of environmental and genetic contributions to DNA methylation and not only their individual contribution.

The main objective of the present study was to further explore these combined effects on DNA methylation at VMRs and their potential functional and disease-related relevance. We investigated the influence of the prenatal environment and genotype on VMRs in the DNA of 2,934 newborns within 4 different cohorts: Prediciton and Prevention of Pre-eclampsia and Intrauterine Growth Restrictions (PREDO, cordblood) ^35^, the UCI cohort (^36-38^, heel prick), the Drakenstein Child Health Study (DCHS, cordblood) ^39; 40^ and the Norwegian Mother and Child Cohort Study (MoBa, cordblood ^41^). Environmental factors in all cohorts were assessed during pregnancy and included metabolic risk factors, hypertension, betamethasone treatment, maternal smoking, demographic factors such as parity and maternal age as well as repeated psychometric evaluations of depression and anxiety.

We focused on the association of these prenatal environmental factors with DNA methylation of VMRs together with genetic variability, exploring both additive (G+E) and interactive (GxE) models, in addition to main effects of genetic (G) and main environmental (E) variables. We also employed the machine-learning algorithm DeepSEA to predict SNP function with regards to regulatory elements ^42^ within the different models. Based on this annotation, we compared the predicted function of both the CpG regions and the SNPs involved in the different models with best prediction by either E, G, G+E or GxE and also tested for enrichment of the relevant SNPs in GWAS both for common psychiatric disorders such as major depressive disorder and schizophrenia and also for type 2 diabetes, inflammatory bowel disease and BMI.

## Results

A general overview of the workflow is depicted in Figure 1.

**Figure 1.**
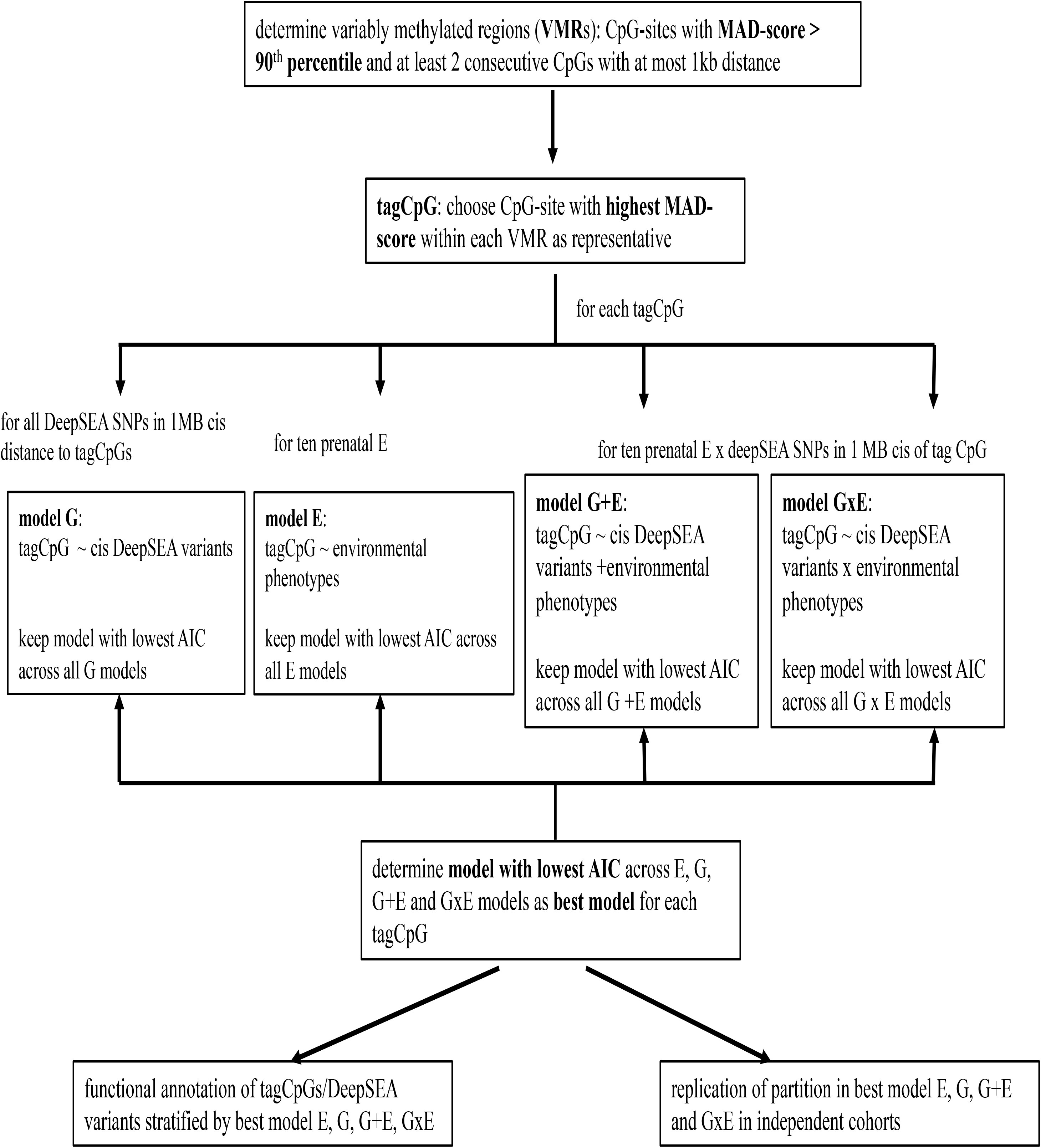
Flow diagram of VMR analysis

### Explaining variance in DNA methylation

We analyzed 963 cord blood samples from the PREDO cohort with available genome-wide DNA methylation and genotype data. Of these samples, 817 had data on the Illumina 450k array (PREDO I) and 146 on the Illumina EPIC array (PREDO II). The main analyses are reported for PREDO I, and replication and extension of the results is shown for PREDO II as well as for three independent cohorts including 121 heel prick samples (UCI cohort, EPIC array) as well as 258 (Drakenstein, 450K and EPIC array) and 1,592 cord blood samples (MoBa, 450K array). Specifically, we tested ten different prenatal environmental factors covering a broad spectrum of prenatal phenotypes (see Table 1) (referred to as “E”), as well as cis SNP genotype (referred to as “G”), i.e., SNPs located in at most 1MB distance to the specific CpG, additive effects of cis SNP genotype and prenatal environment (“G+E”) and cis SNP x environment interactions (“GxE”) for association with DNA methylation levels (see Figure 1). We then assessed for each CpG independently which model described the variance of DNAm best using Akaike’s information criterion (AIC) ^64^. The AIC is based on a goodness-of-fit measure combined with a penalization term for the number of model parameters. In all models, we corrected for child’s gender, ethnicity (using MDS-components) as well as estimated cell proportions to account for cellular heterogeneity.

**Table 1:** overview of investigated cohorts

| cohort | PREDO I | PREDO II | DCHS I | DCHS II | UCI |
| --- | --- | --- | --- | --- | --- |
| sample-size | 817 | 146 | 107 | 151 | 121 |
| methylation array | Illumina 450K | Illumina EPIC | Illumina 450K | Illumina EPIC | Illumina EPIC |
| # VMRs | 3,982 | 8,547 | 6,072 | 10,005 | 9,525 |
| infant gender<br>male | 433 (53.0%) | 75 (51.4%) | 63 (58.8%) | 83 (55.0%) | 65 (53.7%) |
| maternal age mean (sd) | 33.28 (5.79) | 32.25 (4.92) | 26.27 (5.87) | 27.42 (5.93) | 28.47 (4.91) |
| partity mean (sd) | 1,05 (1.02) | 0.87 (1.03) | 0.98 (1.12) | 1.09 (1.07) | 1.11 (1.15) |
| Caesarian section | 169 (20.7%) | 36 (24.7%) | 19 (17.6%) | 35 (23.2%) | 37 (30.6%) |
| pre-pregnancy BMI mean (sd) | 27.42 (6.40) | 25.37 (5.79) | not available | not available | 27.90 (6.44) |
| maternal smoking yes | exclusion<br>criterion | exclusion<br>criterion | 7.40 (10.52) <sup>a)</sup> | 4.94 (9.43) <sup>a)</sup> | 10 (8.2%) |
| gestational diabetes yes | 183 (22.4%) | 19 (13.0%) | no cases<br>available | no cases<br>available | 9 (7.4%) |
| hypertension yes | 275 (33.7%) | 31 (21.2%) | 2 (.19%) | 2 (1.3%) | 7 (5.8%) |
| betamethasone treatment yes | 35 (4.3%) | 2 (1.5%) | not available | not available | no cases<br>available |
| anxiety score mean (sd) | 33.93 (7.90) <sup>b)</sup> | 34.43 (8.38) <sup>b)</sup> | 5.70 (4.15) <sup>c)</sup> | 5.32 (3.91) <sup>c)</sup> | 1.67 (0.41) <sup>d)</sup> |
| depression score mean (sd) | 11.34 (6.47) <sup>e)</sup> | 11.53 (6.98) <sup>e)</sup> | 17.64 (12.10) <sup>f)</sup> | 12.52 (11.55) <sup>f)</sup> | 0.68 (0.41) <sup>g)</sup> |
| proportion:<br>best model E | 2.0% | <1% | <1% | <1% | 4.1% |
| best model G | 45.1% | 15.0% | 15.8% | 11.5% | 12.8% |
| best model G+E | 23.9% | 29.0% | 29.8% | 32.1% | 24.1% |
| best model GxE | 29.0% | 56.0% | 54.3% | 56.3% | 59.0% |

<sup>a)</sup> based on ASSIST Tobacco Score<sup>b)</sup> STAI sum scores<sup>c)</sup> SRQ-20<sup>d)</sup> STAI average scores<sup>e)</sup> CESD sum scores<sup>f)</sup> BDI-II<sup>g)</sup> CESD average score

### Variably Methylated Regions

We analyzed VMRs, defined as regions of CpG-sites showing the highest variability across all methylation sites. In PREDO I, we identified 10,452 variable CpGs that clustered into 3,982 VMRs (see Table S1).

While overall methylation levels at CpGs are bimodally distributed with peaks at very low methylation (beta-value < 0.1) and very high methylation (beta-value > 0.8), the distribution of methylation at CpGs in VMRs is unimodal and VMRs presented with intermediate methylation levels (median beta-value=0.52, 25^th^-75^th^ percentile = 0.39-0.62, see Figure 2A). As compared to all CpG-sites on the 450K array, VMRs in cord blood were enriched for OpenSea (p=2.70 × 10^−03^, Fisher-test) and Shores (p=9.39 × 10^−91^) and depleted for Islands (p=2.35 × 10^−32^). Furthermore, VMRs were enriched for distal intergenic regions (p=2.19 × 10^−31^, Fisher-test) and depleted for introns (p=1.26 × 10^−07^). Additionally, we checked if VMRs were enriched for transcription factor (TF) binding sites. Using the data of ReMap ^65^ and ENCODE ^66^ we found significant enrichment for VMRs (p=6.25 × 10^−61^, Fisher-test) as compared to non-VMRs on the 450K array. Here, 82% of all VMR-CpGs overlapped with at least one TF, with CTCF and NR3C3 being among the top enriched factors.

**Figure 2.**
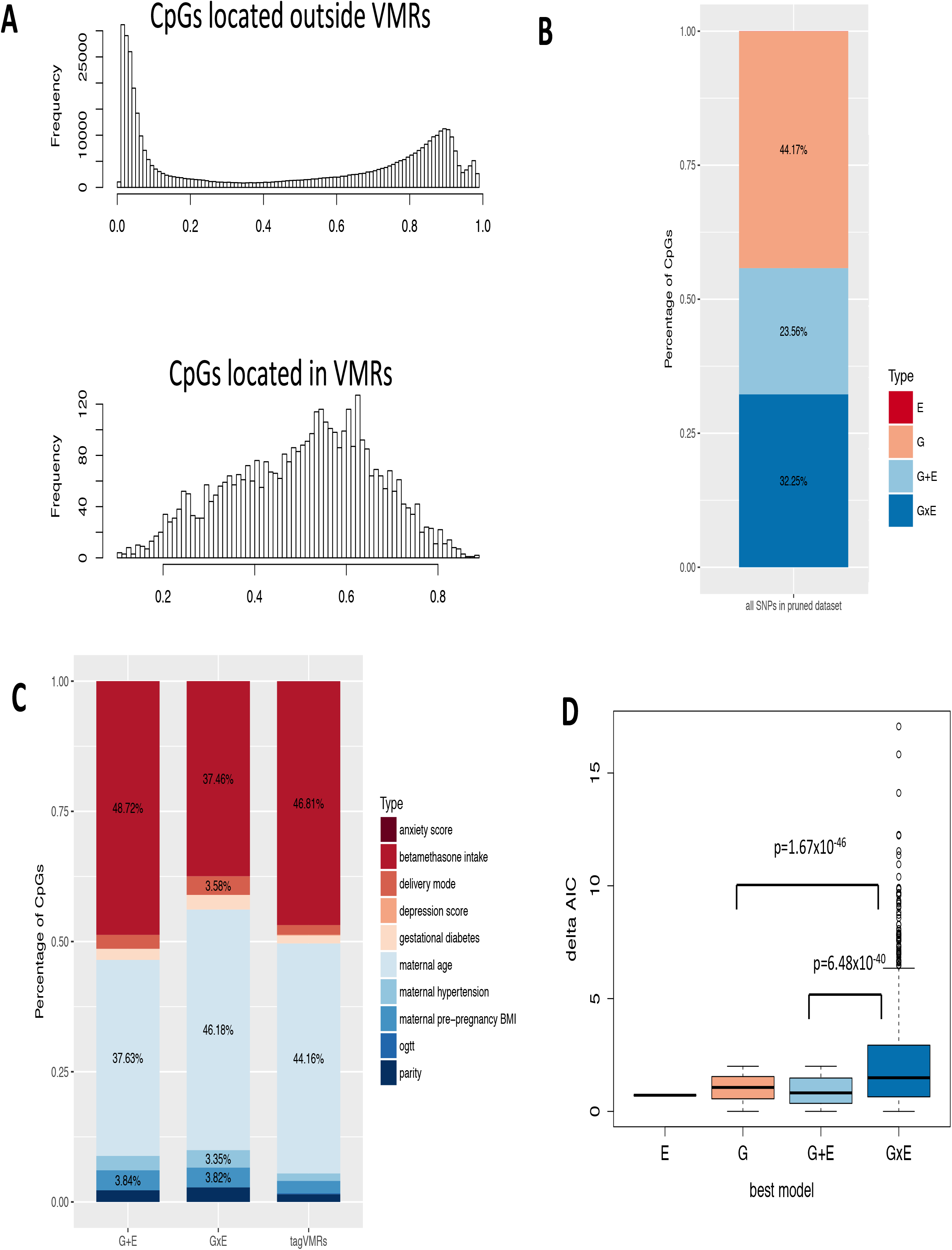
A: Histogram of median methylation levels of CpG-sites located in non-VMRs (above) and CpG-sites located in VMRs (below). B: Percentage of models (G, E, GxE or G+E) with the lowest AIC explaining variable DNA methylation using the PREDO I dataset with pruned SNPs. C: Distribution of the different types of prenatal environment included in the E model with the lowest AIC (right), in the combinations yielding the best model GxE (middle), or the best model G+E models (left). To increase readability all counts < 3% have been omitted. D: DeltaAIC, i.e, the difference in AIC, between best model and next best model, stratified by the best model. Y-axis denotes the delta AIC and the X-axis the different models. The median is depicted by a black line, the rectangle spans the first quartile to the third quartile, whiskers above and below the box show the location of minimum and maximum beta-values.

VMRs have been associated with specific chromatin states ^67^. As compared to non-VMRs, VMRs in our dataset were depleted for active and flanking transcription start sites (TSS), for strong transcription and for transcription at 5’ and 3’. In contrast, VMRs were enriched for weak transcription, enhancers, ZNF genes and repeats, heterochromatin, bivalent/poised TSS, bivalent flanking TSS/enhancers, bivalent enhancers, repressed and weak repressed PolyComb sites (Figure S1).

We used the publicly available eQTM results from Bonder et al. ^68^, who examined whether VMRs significantly overlapped with expression quantitative trait methylation sites (eQTMs), i.e., CpGs significantly associated with gene expression. As the analysis presented by Bonder et al. was based on CpGs located in proximity to the TSS of the specific transcript, we used only PREDO I VMRs also located within genes (n=5,905). The overlap between significantly associated eQTMs from Bonder et al. and VMRs in PREDO I was significantly higher than expected by chance (p=9.99 × 10^−05^, based on sampling of 10,000 random CpG-sets), revealing that areas with high levels of inter-individual variation in DNA methylation overlap strongly with sites associated with gene expression.

Additionally, 6,074 CpG-sites significantly associated with maternal smoking ^11^ and 104 CpG-sites significantly associated with maternal BMI ^13^ significantly overlapped (p=0.009 for smoking and p=0.0009 for BMI, based on sampling of 1,000 random CpGs-sets) with VMR CpGs. Furthermore, CpG-sites reported in ^11; 13^ showed higher MAD-scores in PREDO I as compared to CpGs which were not significantly associated in either of these studies.

Our results confirm those of previous studies showing VMRs are enriched in specific functional regions of the genome, correlate with differences in gene expression, and overlap with sites associated with specific prenatal environmental factors.

We next examined the factors that best explained the variance in methylation in these functionally relevant sites. For each VMR, we chose the CpG-site with the highest MAD-score as representative of the whole region. These CpGs are named tagCpGs.

### G, E, G+E or GxE?

We compared the fit of four models for each of the 3,982 tagCpGs (see Figure 1): best SNP (G model), best environment (E model), SNP + environment (G+E model) and SNP x environment (GxE model). Top association results for each model are depicted in Tables S2-S5. For each tagCpG, the model with the lowest AIC was chosen as the best model (see Methods section). In total, 44 % of tagCpGs were best explained by G (n=1,759), followed by GxE (32%, n=1, 284) and G+E (24%, n=938) (Figure 2B). E explained most variance in one tag CpG. All tag CpGs and the respective SNPs and environments from the best model are listed in Tables S6-S9.

With regard to environmental factors, 48.7% of tagCpGs best explained by the G+E model were associated with environmental factors related with stress or, in particular, glucocorticoids (i.e., maternal betamethasone treatment), 42.5% with general maternal factors (mostly maternal age) and 8.8% with factors related to metabolism (pre-pregnancy BMI, hypertension). For best model GxE, the proportions were similar with 37.4%, 52.6% and 10%, respectively (see Figure 2C).

Finally, we looked into the delta AIC, i.e., the difference between the AIC of the best model to the AIC of the next best model. If the best model was G (n=1,759), the next best model was G+E in 75% of the cases (n=1,313), and GxE for the remaining part (n=446). For the 1,248 tagCpGs where the best model was GxE, the next best model was mostly G (n=747) followed by the additive model (n=537). In the case of the 938 tag CpGs with best model G+E, the next best model was mostly GxE (n=527), followed by G only (n=411).

Interestingly, E never occurred as next best model. For the one CpG with best model E, the next best model was G. The delta AIC for best model GxE to the next best model was significantly higher (mean delta AIC=2.16) as compared to CpGs with G as the best model (mean delta AIC=1.04, p= 6.48 × 10^−40^, Wilcoxon-test) or G+E as the best model (mean delta AIC=0.93, p=1.67 × 10^−46^, Figure 2D). Furthermore, the delta AIC for best model G (mean=1.04) was significantly higher as compared to best model G+E (mean=0.93, p=2.56 × 10^−06^).

We conclude that GxE models are selected by larger margins while best G and best G+E models are selected with lower margins.

### Functional mapping – DeepSEA prediction

The results presented so far are based on linkage disequilibrium (LD)-pruned SNP data, so it is not necessarily the functional SNP but any tag SNP that was likely selected in the models. Due to the nature of LD-structure, a functional SNP with a high correlation to the representative SNP can equally be favored for analysis. To better understand the contribution of functional SNPs to VMRs we restricted the analyses only to potentially functional relevant SNPs using DeepSEA ^42^. DeepSEA, a deep neural network pretrained with DNase-seq and ChIP-seq data from the ENCODE ^66^ project, predicts the presence of histone marks, DNase hypersensitive regions (DHS) or TF binding for a given 1kb sequence. The likelihood that a specific genetic variant influences regulatory chromatin features is estimated by comparing predicted probabilities of two sequences where the bases at the central position are the reference and alternative alleles of a given variant. We reran the four models restricting the cis-SNPs to those 36,241 predicted DeepSEA variants that were available in our imputed, quality-controlled genotype dataset.

Top results for models including G, GxE and G+E are depicted in Tables S10-S12.

Results were comparable to what we observed before: 1,796 (45.2%) of tagCpGs presented with best model G, 948 CpGs (23.9%) with best model G+E, 1,151 CpGs (24 %) with best model GxE and 77 CpGs (2%) with best model E (Figure 3A). Only 10 tagCpGs did not present with any DeepSEA variant within 1MB distance in cis and were therefore not considered further. All respective CpG-environment-DeepSea SNP-combinations are depicted in Tables S13-S16.

**Figure 3.**
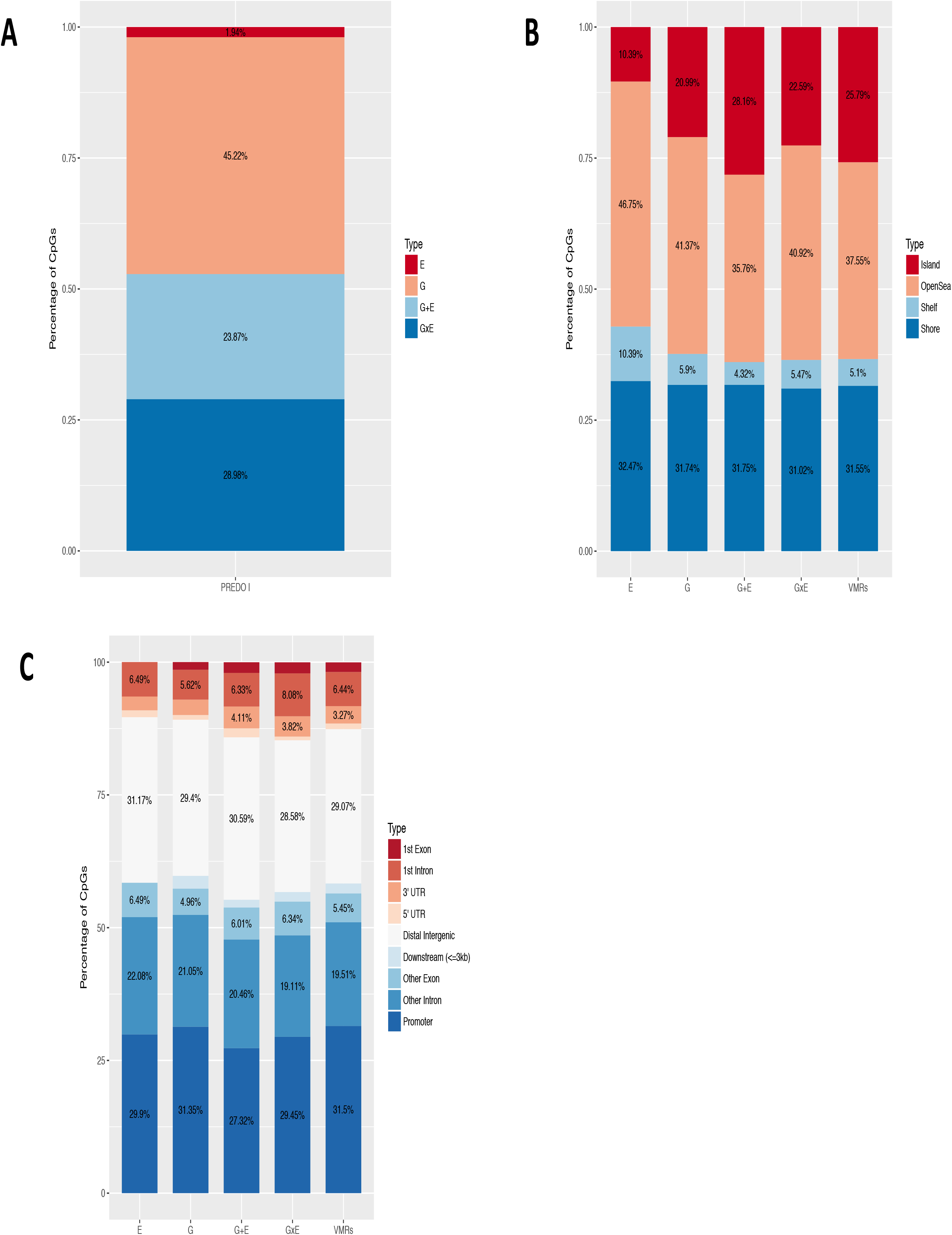
A: Percentage of models (G, E, GxE or G+E) with the lowest AIC explaining variable DNA methylation using the dataset with DeepSEA annotated SNPs. B: Distribution of the locations of all VMRs and tagVMRs with best model E, G, G+E and GxE on the 450k array using only DeepSEA variants in relationship to CpG-Islands based on the Illumina 450K annotation. C: Distribution of gene-centric locations of all VMRs and tagVMRs with best model E, G, G+E and GxE on the 450k array using only DeepSEA variants.

In conclusion, also when we focus on functionally relevant SNPs, it is mainly the combination of genotype and environment which explains VMRs best.

### Are VMRs best explained by E alone, G alone, G+E or by GxE different from each other?

Focusing on combinations between tagCpGs, environmental factors and DeepSEA variants, we functionally annotated both the SNPs as well as the tagCpGs within the different models. TagCpGs best explained by G and GxE were enriched for OpenSea (p=5.16 × 10^−03^), G+E tagCpGs were enriched for Islands (p= 1.04 × 10^−05^) and tagCpGs best explained by E were depleted for Islands (p= 3.10 × 10^−03^, Figure 3B). Furthermore, GxE and G+E tagCpGs were depleted for promoters (p=2.95 × 10^−02^, Figure 3C).

With regard to enrichment/depletion of specific histone marks based on the ENCODE data ^66^ in comparison to all VMRs, E tagCpGs were nominal significantly depleted for active TSS (p=0.03), for bivalent enhancers (p=0.02) and repressed PolyComb (p=0.001).

GxE tagCpGs were significantly depleted for flanking active TSS (p=0.02) and for flanking bivalent TSS/enhancers (p=0.006). G+E tagCpGs were significantly enriched for flanking bivalent TSS/enhancers (p=0.009), for bivalent enhancers (p=0.03) and for bivalent/poised TSS (p=0.004) and depleted for genic enhancers (p=0.03, see Figure 4A and 4B).

**Figure 4.**
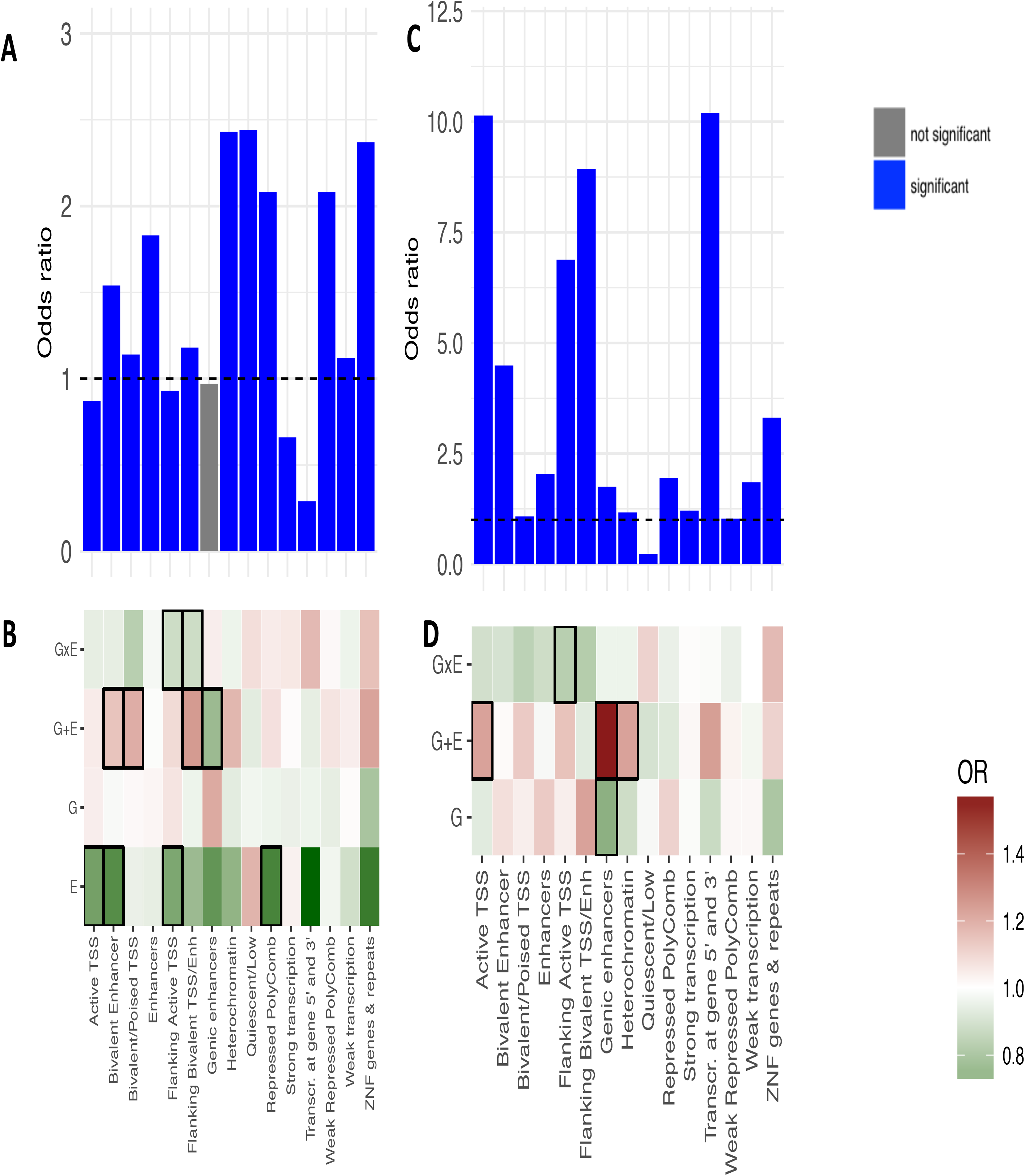
A: Histone mark enrichment for all VMRs. The Y-axis denotes the fold enrichment/depletion as compared to no-VMRs. Blue bars indicate significant enrichment/depletion, grey bars non-significant differences. B: Histone mark enrichment for tagVMRs with best model E, G, G+E and GxE relative to all VMRs. Green color indicates depletion, red color indicates enrichment. Thick black lines around the rectangles indicate significant enrichment/depletion. C: Histone mark enrichment for all DeepSEA variants in the dataset. Blue bars indicate significant enrichment/depletion. D: Histone mark enrichment for all DeepSEA variants involved in models where either G, G+E or GxE is the best model as compared to all tested DeepSEA variants. Green color indicates depletion, red color indicates enrichment. Thick black lines around the rectangles indicate significant enrichment/depletion.

Overall, tagCpGs with best model G, G+E or GxE were located in functionally distinct regions, with GxE tagCpGs being enriched for OpenSea positions and depleted for TSS markers.

### Are functional SNPs which are only involved in G models different from functional SNPs only involved in G+E/GxE models?

Overall, 1,387 DeepSEA variants uniquely involved in best G models, 867 were uniquely in best GxE models and 706 uniquely in best G+E models. As a DeepSEA variant can be in multiple 1 MB-cis windows around the tagCpGs, several DeepSEA variants were involved in multiple best models: 147 DeepSEA variants overlapped between G and GxE, 137 between G and G+E and 94 between GxE and G+E VMRs. We observed no significant differences with regard to gene-centric location for DeepSEA variants involved only in G models, only in GxE models or in multiple models. However, DeepSEA variants involved only in G+E models were significantly enriched for promoter locations (p=1.06 × 10^−03^, see Figure S2A).

Although no significant differences were present, DeepSEA SNPs involved in the G and G+E model were located in closer proximity to the specific CpG (model G: mean absolute distance=259.6 kb, sd=289.1 kb, model G+E: mean absolute distance =250.5 kb, sd=283.9 kb, Figure S2B) whereas DeepSEA SNPs involved in GxE models (mean absolute distance =363.9 kb, sd=309.3 kb) showed broader peaks around the CpGs.

We next evaluated if functionally relevant G, GxE and G+E SNPs differed with regard to enrichment for histone marks. As expected, DeepSEA variants in general were enriched across multiple histone marks indicative of active transcriptional regulation (Figure 4C). DeepSEA variants involved in best model G+E showed further enrichment for TSS marks as compared to all tested DeepSEA variants. In contrast, GxE DeepSEA variants were significantly depleted in such regions. G+E SNPs, but not the other SNP types were highly enriched for genic enhancers and heterochromatin while G SNPs were depleted for genic enhancers (Figure 4D).

### Is the proportion of best models dependent on the variability of CpG-sites?

The power to detect meQTLs depends on the variance of DNA methylation at the specific CpG-site ^69^ and the allele frequency of SNPs mapped in close proximity to the most variably methylated sites ^70^. Therefore, VMRs, which by definition are restricted to CpG-sites with a high variability, might not only be biased for the identification of significant meQTLs but also have a higher proportion of the variance explained by G using the AIC. We investigated if this was true by re-running the E, G, G+E and GxE models on all CpGs-sites, regardless of whether they were located in VMRs or not. As maternal age was one of the most important predictors for methylation levels at VMRs in our analysis (see Figure 2C), we focused on this phenotype. We found that, across all CpG-sites and all variability levels (see Figure S3), the pattern of best models remained stable indicating that, at least in our sample, combined G and E effects are also present in sites not located in VMRs and in less variable sites.

### Replication of relative distribution of best models in independent cohorts

To assess whether the relative distribution of the best models for VMRs and DeepSEA variants was stable, we assessed the relative distribution of these models in 3 additional samples with VMR data both from the Illumina 450K as well as the IlluminaHumanEPIC arrays. These included the Drakenstein Child Health Study (DCHS), with DNA methylation assessed by the 450K array in 107 newborns (DCHS I) and with the EPIC array in 151 newborns (DCHS II). The EPIC array was also used in the UCI cohort with 121 neonates as well as in PREDO II with 146 neonates. While major maternal factors overlapped among the cohorts - such as maternal age, delivery method, parity and depression during pregnancy - there were also differences, as the non-PREDO cohorts did not include betamethasone treatment but did include maternal smoking (see Table 1). VMRs differed between cohorts: overall 6,251 CpGs assessed on the 450K array were located within VMRs of PREDO I as well as in the VMRs of DCHS I. With regard to the EPIC array: 4,634 CpGs were located within VMRs of PREDO II as well as in the VMRs of DCHS II and the UCI cohort.

Nevertheless, the overall pattern remained stable: in all 4 analyses, DCHS I, DCHS II, UCI and PREDO II, we replicated that E alone models almost never explained most of the variances, while G alone models explained the most variance in up to 15% of the VMRs; G+E in up to 29%; and GxE models in up to 59% (see Figure 5 and Table 1). We were also able to replicate our finding showing that GxE VMRs were enriched for OpenSea positions on the 450K array (DCHS I, OR=1.11, p= 3.50 × 10^−02^). This remained true in data from the EPIC array, which in itself contains more CpGs in the OpenSea regions as compared to the 450k as GxE were enriched in OpenSea regions over the EPIC content (PREDO II: OR=1.23, p=1.19 × 10^−06^, DCHS II: OR=1.13, p=1.00 × 10^−04^). For all cohorts, the delta AIC for best model GxE to the next best model was also significantly higher as compared to CpGs with G, E or G+E as the best model. The additional cohorts thus showed a consistent replication of the fact the GxE and G+E models best explained the variance of the majority of the VMRs; the VMRs explained by a combination of G and E were enriched in OpenSea regions; and that GxE models were always selected by a higher margin than other models.

**Figure 5.**
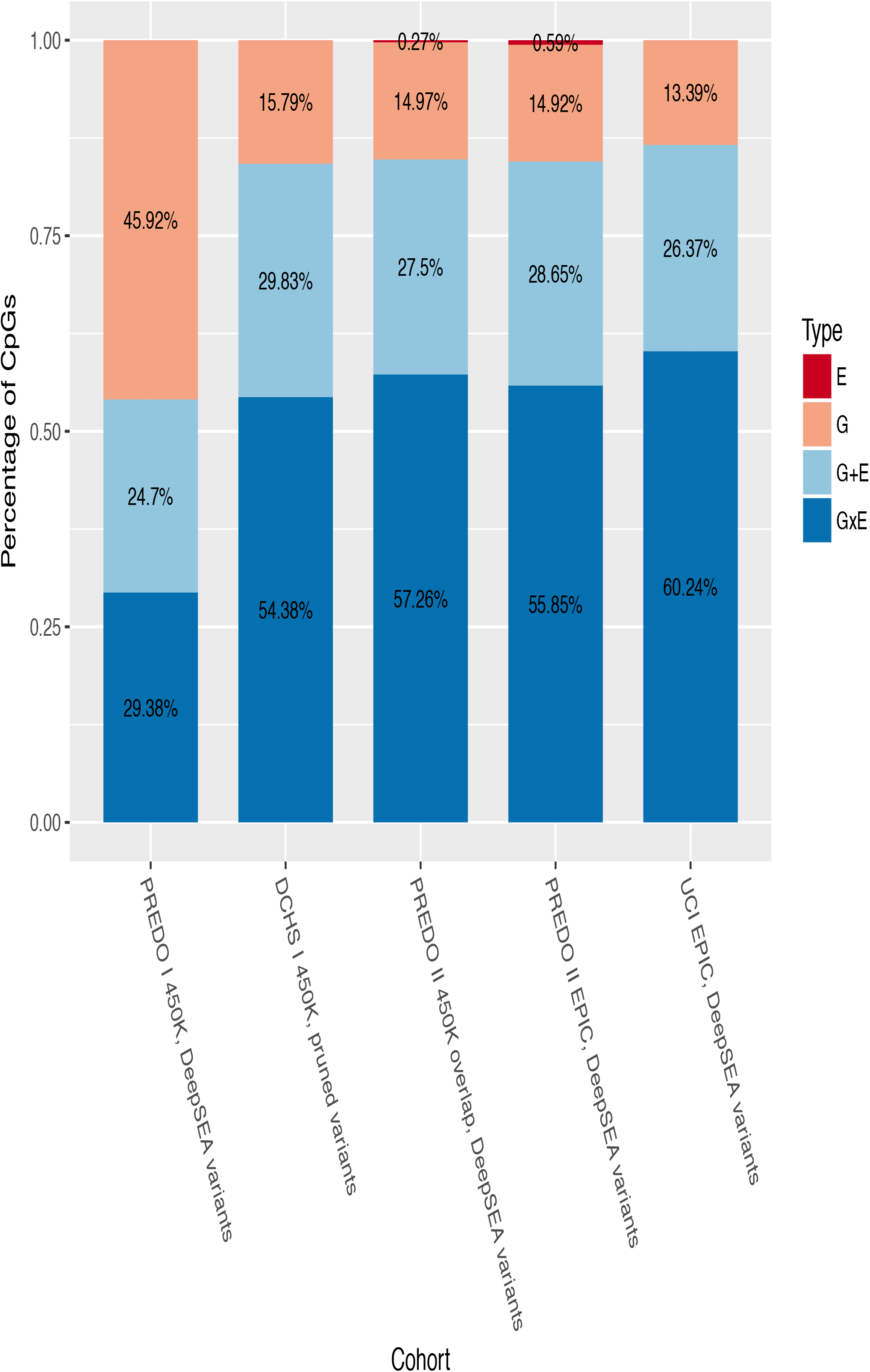
Percentage of models (G, E, GxE or G+E) with the lowest AIC explaining variable DNA methylation in PREDO I (450K), DCHS I (450K), PREDO II (EPIC), MecTransGen (EPIC) and DCHS II (EPIC)

### Association with smoking

As we did not observe significant main E effects on DNA methylation for most of the tested Es in our cohorts, we chose to rerun the analyses focusing on maternal smoking, described as one of the most highly replicated factors shaping the newborns’ methylome ^11^. This would allow an assessment of how inclusion of a validated E factor would influence the relative distribution of the best models. We first ran traditional epigenome-wide association analyses for maternal smoking in the cohorts where this exposure was included, namely the UCI, DCHS I and DCHS II. We observed that those CpG-sites where association with smoking was nominally significant in our samples, were significantly enriched for CpG-sites which had been reported to be associated with this exposure in the meta-analysis by Joubert et al. ^11^ at an FDR corrected p-value cut-off of 0.05 (UCI, p=1.30 × 10^−28^, OR=1.80, DCHS I, p=2.77 × 10^−32^, OR=1.80, DCHS II p=1.35 × 10^−15^, OR=1.56). Next, we tested whether those CpGs that were associated with maternal smoking in Joubert et al. ^11^ at FDR 0.05, presented with best model E=smoking, or if the inclusion of genotype yielded a better model. For UCI 5,362 CpGs out of the 6,073 reported CpGs were available. From these, 25 (<1 %) were best explained by smoking alone (E), whereas 5,001 (93.3%) were best explained by genotype (G), 139 (2.6%) by G+maternal smoking (G+E) and 197 (3.6%) by Gxmaternal smoking (GxE). In DCHS I, 5633 of the top CpGs were available, 4,773 (84.7%) presented with best model G, 603 (10.7%) with best model GxE and 257 (4.6 %) with best model G+E. In DCHS II, out of 5,405 CpGs, 2 (< 1%) presented with best model E, 3,058 (56.6%) with best model G, 1,654 (30.6%) with best model GxE and 691 (12.8%) with best model G+E. This underscores the point that even for phenotypes, such as maternal smoking, with documented main E effects on cord blood methylation, genotypic information should be considered.

### Replication of GxE and G+E findings

We focused on choosing the best model according to the AIC. Although we are likely underpowered to robustly detect significant GxE interactions, we examined if combined effects of genotype and environment in the CpGs that were identified to be best explained by G+E and GxE in PREDO I, could be replicated in an independent sample. We evaluated whether specific G+E and GxE effects on cord blood methylation measured using the Illumina 450K array were also present in the MoBa cohort including 1,592 newborns. We restricted the analysis to those combinations of CpG-DeepSEA SNP and environments that were nominally significant for GxE or G+E and also presented with the lowest AIC for this specific model in PREDO I. Overall, 913 GxE and 372 G+E combinations fulfilled this criterion. Of these, 515 GxE and 178 G+E combinations were also available in the MoBa cohort. Overall, 268 GxE and 98 G+E combinations showed the same direction of effects in both cohorts and we found no significant difference between the direction of effects in both cohorts, neither for GxE (p=0.176, sign-test) nor for G+E combinations (p=0.88). Overall, 9 GxE and 15 G+E combinations were nominally significant in MoBa and showed the same direction of effects as in PREDO I. Next, we combined both studies via random-effects-meta-analyses where 6 GxE and 18 G+E combinations were significantly associated and survived multiple testing across all tested combinations (FDR 0.05), showing the same direction of effects in both cohorts and presenting with lower p-values in the meta-analysis as compared to PREDO I alone. These results are depicted in Table S17 and S18. Two of the tophits for GxE and G+E are presented in Figure 6A and B, respectively.

**Figure 6.**
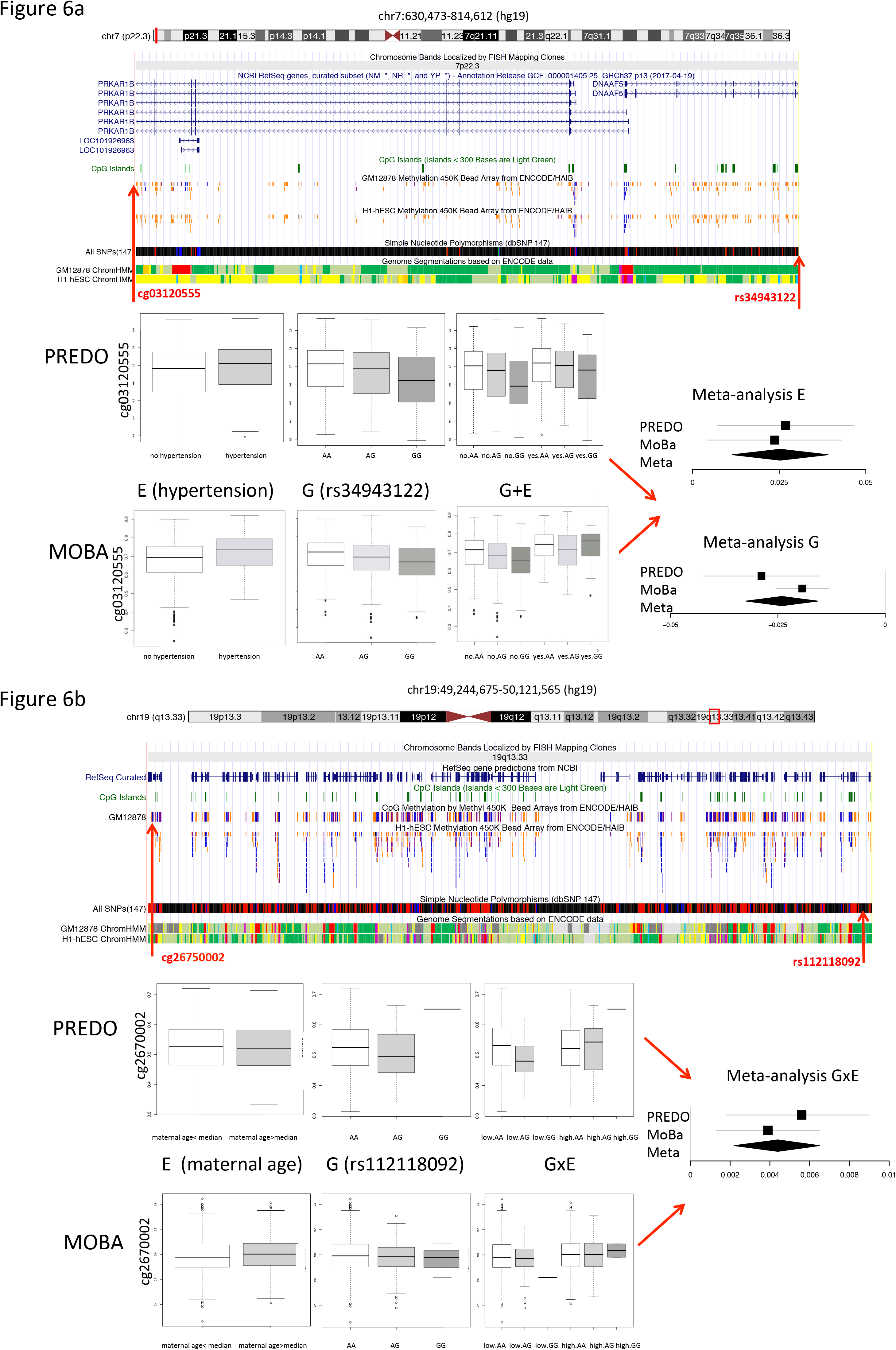
A: Illustration of one of the top G+E hits from the meta-analysis, including location of SNP rs34943122 and CpG cg03120555 relative to their genomic location and the following UCSC tracks (based on gh19): NCBI RefSeq genes, CpG Islands, GM12878 Methylation 450k BeadArray from ENCODE/HAIB, H1-hESC Methylation 450k BeadArray from ENCODE/HAIB, simple nucleotide polymorphism (db SNP147), genome segmentations based on ENCODE data for GM12878, genome segmentations based on ENCODE data for H1-hESC. In the lower part of the panel, boxplots are given for the results from PREDO (upper panels) and MoBa (lower panels) for main effects of E (hypertension no-yes), G and G+E). The Y-axis denotes the respective beta-values and the X-axis the different environmental conditions or genotypes. The median is depicted by a black line, the rectangle spans the first quartile to the third quartile, whiskers above and below the box show the location of minimum and maximum beta-values. On the right side, a forest-plot for this G+E combination is given where the effect size estimate is depicted as black square and the grey line indicates the respective confidence interval on the X-axis. The Y-axis denotes the different studies and the meta-analysis. The result of the meta-analysis is depicted as a diamond: the center line of the diamond gives the effect size estimator from the meta-analysis while the lateral tips of the diamond indicated the lower and upper limits of the confidence interval. B: Illustration of one of the top GxE hits from the meta-analysis, including location of SNP rs112118092 and CpG cg26750002 relative to their genomic location and the following UCSC tracks (based on hg19): NCBI RefSeq genes, CpG Islands, GM12878 Methylation 450k BeadArray from ENCODE/HAIB, H1-hESC Methylation 450k BeadArray from ENCODE/HAIB, simple nucleotide polymorphism (db SNP147), genome segmentations based on ENCODE data for GM12878, genome segmentations based on ENCODE data for H1-hESC. In the lower part of the panel, boxplots are given for the results from PREDO (upper panels) and MoBa (lower panels) for main effects of E (maternal age below the median, maternal age above the median), G and GxE. The Y-axis denotes the respective beta-values and the X-axis the different environmental conditions or genotypes. The median is depicted by a black line, the rectangle spans the first quartile to the third quartile, whisker above and below the box show the location of minimum and maximum beta-values. On the right side, a forest-plot for this GxE combination is given where the effect size estimate is depicted as black square and the grey line indicates the respective confidence interval on the X-axis. The Y-axis denotes the different studies and the meta-analysis The result of the meta-analysis is depicted as a diamond: the center line of the diamond gives the effect size estimator from the meta-analysis while the lateral tips of the diamond indicated the lower and upper limits of the confidence interval.

### Disease relevance

Finally, we tested whether functional DeepSEA SNPs involved in only G, only GxE and only G+E models in PREDO I for their enrichment in GWAS hits. We used all functional SNPs and their LD proxies (defined as r^2^ of at least 0.8 in the PREDO cohort and in maximal distance of 1MB to the target SNP) and performed enrichment analysis with the overlap of nominal significant GWAS hits. We selected for a broad spectrum of GWAS, including GWAS for complex disorders for which differences in prenatal environment are established as risk factors, but also including GWAS on other complex diseases. For psychiatric disorders, we used summary statistics of the Psychiatric Genomics Consortium (PGC) including association studies for autism ^71^, attention-deficit-hyperactivity disorder ^72^, bipolar disorder ^73^, major depressive disorder ^74^, schizophrenia ^75^ and the cross-disorder associations including all five of these disorders ^76^. Additionally, we included GWAS of inflammatory bowel disease ^77^, type 2 diabetes ^78^ and for BMI ^79^. Nominal significant GWAS findings were enriched for DeepSEA variants and their LD proxies per se across psychiatric as well as non-psychiatric diseases (Figure 7A). However, G, GxE and G+E DeepSEA variants showed a differential enrichment pattern above all DeepSEA variants (Figure 7B), with strongest enrichments of GxE DeepSEA variants in GWAS of autism spectrum disorder (p=1.32 × 10^−12^,OR=1.16 above DeepSEA), bipolar disorder (p=4.90 × 10^−06^, OR=1.16) and major depressive disorder (p=1.88 × 10^−26^, OR=1.16) and G+E DeepSEA variants in GWAS for schizophrenia (p=2.41 × 10^−58^, OR=1.27) and inflammatory bowel disease (p=9.14 × 10^−124^, OR=1.52). While SNPs with strong main meQTL effects such as those within G and G+E models have been reported to be enriched in GWAS for common disease, we now also show this for SNPs within GxE models that often have non-significant main G effects.

**Figure 7.**
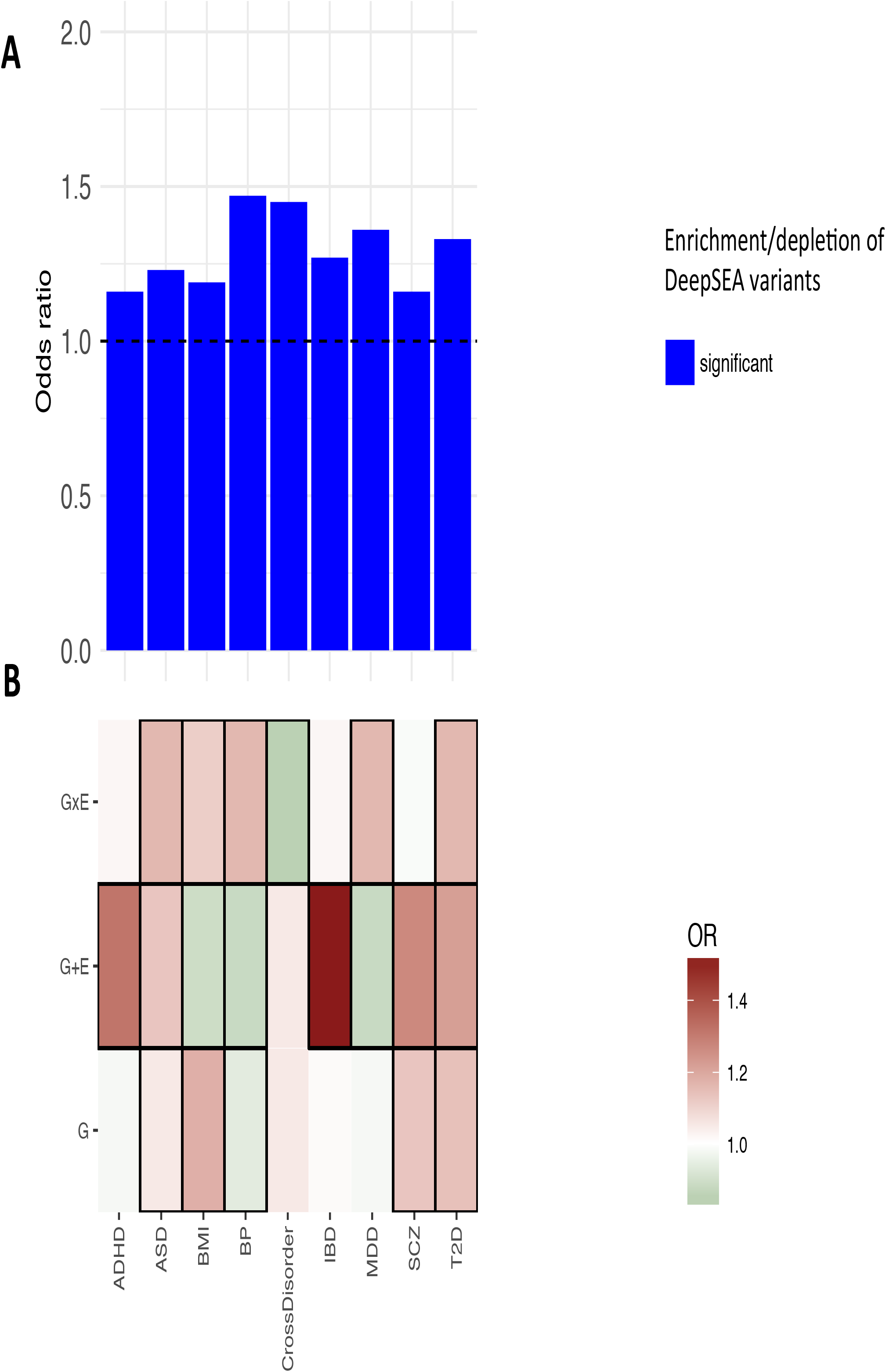
A: Enrichment for nominal significant GWAS associations for all tested DeepSEA variants and their LD proxies for GWAS for ADHD (attention deficit hyperactivity disorder), ASD (autism spectrum disorder), BMI (body-mass index), BP (bipolar disorder), CrossDisorder, IBD (inflammatory bowel disease), MDD (major depressive disorder), SCZ (schizophrenia) and T2D (Type 2 diabetes). The Y-axis denotes the fold enrichment non-DeepSEAvariants. Blue bars indicate significant enrichment/depletion. B: Enrichment for nominal significant GWAS hits for DeepSEA variants and their LD proxies involved in best models with G, G+E or GxE as compared to all tested DeepSEA variants. Green color indicates depletion, red color indicates enrichment. Thick black lines around the rectangles indicate significant enrichment/depletion.

## Discussion

We evaluated the effects of prenatal environmental factors and genotype on DNA methylation at VMRs identified in neonatal blood samples. We found that most variable methylation sites were best explained by either genotype and prenatal environment interactions (GxE) or additive effects (G+E) of these factors, followed by main genotype effects. This pattern was replicated in independent cohorts and underscores the need to consider genotype in the study of environmental effects on DNA methylation. This pattern held true when specifically examining the E of maternal smoking, which has the most robust environmental influence on neonatal DNA methylation established to date ^11^. We observed that DNA methylation of CpG sites associated with maternal smoking were also best explained by a combination of G and E and not E alone in our cohorts. This finding is in agreement with a previous study of Gonseth et al. ^33^ who reported significant genetic contribution to CpG-sites that are sensitive to maternal smoking and hence advocated an adequate inclusion of genotype information in DNA methylation studies. Our results support that a more complete picture of factors influencing DNA methylation should include genotype information and could also help to detect environmental influences that may only be unmasked in the context of a specific genetic background.

VMRs best explained by G, G+E or GxE and their associated functional genetic variants were located in distinct genomic regions, suggesting that different combinatorial effects of G and E may impact VMRs with distinct downstream regulatory effects. We also observed that functional variants with best models G, G+E or GxE, all showed significant enrichment within GWAS signals for complex disorders beyond the enrichment of the functional variants themselves. While this was expected for G and G+E models based on results from previous studies ^24; 26; 27; 29^, it was surprising for GxE SNPs, as these often do not have highly significant main genetic effects. Their specific enrichment in GWAS for common disorders supports the importance of these genetic variants that moderate environmental impact both at the level of DNA methylation but also, potentially, for disease risk.

Interestingly, the best models with G+E or GxE, i.e. with the lowest AIC, were not necessarily the combination of the SNP with the lowest AIC with the environment with the lowest AIC. While nearly 80% of all G+E models contained these specific SNP and environment combinations, this was only the case for half of the GxE models. This suggests that testing only variables with significant main effects in multiplicative interaction or additive models might be too restrictive and could lead to important combinations of genotype and phenotype not being captured.

The fact that GxE and G+E best explained the majority of VMRs (see Figure 5) and that GxE models were selected by a larger margin than the other models (see Figure 2D) was consistently found across all tested cohorts. These findings are in line with a previous report by Teh et al. ^34^ who performed a similar analysis based on AIC in umbilical cord tissue and also support the hypothesis that E alone was only rarely the best model to explain variance in DNA methylation as well as that GxE models were selected by a larger margin as compared to G models. In the Teh et al study, 75% of all VMRs were best explained by GxE (the rest by G alone), while in our results this number was only 32%. The most important difference between these two studies is that we additionally tested for additive models and that we took all combinations of cis SNP and environmental factors into account whereas Teh et al. focused on SNPs and phenotypes that presented with lowest AIC values in only G and only E models. Apart from that, several other factors might explain this discrepancy. First, our sample was larger and also ethnically more homogenous as compared to Teh at al. ^34^ who studied a mixed Asian population. Second, not all of the environmental factors that were tested in Teh et al. were available for our cohort. Third, we used imputed SNP genotypes whereas Teh et al. used only measured genotypes. In fact, 64% of the SNPs that are involved in best G models in our dataset were imputed. Fourth, as compared to Teh et al., we looked into a different set of CpGs as VMRs between the two studies only showed a slight overlap (n = 219 tagCpGs), likely due to differences in tissue and ethnicity.

In addition to consistent findings using AIC-based approaches, we also saw some indication for replication of individual GxE and G+E combinations on selected VMRs using p-value based criteria, although we are underpowered to conduct a definitive analysis on this. Using data from PREDO and the even larger MoBa cohort, we observed that the direction of effects for the top G+E and GxE models in PREDO were similar in MoBa, indicating the same directionality of effects across cohorts. This supports the possibility that our findings may be robust not only at the level of AIC for the best models, but also for specific combinations of G and E on VMRs.

With the exclusion of maternal smoking as it was not present in the PREDO cohort, the environmental factors that were most included in the top G+E and GxE models were betamethasone treatment (in 49% of all best G+E and in 38% of all best GxE models) and maternal age (in 36% of all best G+E and in 47% of all best GxE models). It has previously been shown that exogenous as well as endogenous glucocorticoids are associated with differences in DNA methylation with evidence from animal, human and in vitro studies ^31; 80-83^. Effects of maternal age on newborn DNA methylation that may last into adulthood have already been reported and included data from the here used MoBa cohort ^84^. Also in line with smaller or inconsistent effects on DNA methylation reported for maternal pregnancy stress ^15^, prenatal maternal anxiety or depression were not among the stronger environmental factors, in comparison with the other assessed environments that included pharmacologic and metabolic factors.

While E alone was rarely the best model, it should be pointed out that main environmental effects on DNA methylation were observed (see Table S3), and consistent with previous large meta-analyses in the case of maternal smoking. However, the inclusion of genotypic effects in addition explained more of the variance. This supports the view that while main E effects on the newborn methylome are present, genotype is an important factor that, in combination with E, may explain even more of the variance in DNA methylation.

VMRs best explained by either E, G, G+E or GxE and their associated functional SNPs were enriched for distinct genomics locations and chromatin states (see Figure 4). Overall, VMRs best explained by GxE were consistently enriched for regions annotated to the OpenSea, both against the VMRs derived from the Illumina 450K array as well as the EPIC array, the later mainly adding content in OpenSea regions compared to the older array ^85; 86^. OpenSea regions describe regions that have lower CpG density and are located farthest from CpG Islands ^87^. Open Sea regions have been reported to be enriched for environmentally-associated CpGs with for example exposure to childhood trauma ^88^ and may harbor more long-range enhancers.

In addition to their position relative to CpG islands and their CpG content, G, GxE and G+E VMRs and their associated functional SNPs also showed distinct enrichments for chromatin marks. Compared to 450K VMRs in general, that are enriched for enhancers (bivalent and not), VMRs with GxE as the best models were relatively depleted in regions surrounding the TSS, while VMRs with G+E were relatively enriched in these regions (see Figure 4), suggesting that GxE VMRs are located at more distance from the TSS than G+E and G VMRs. To better map the potential functional variants in these models and to compare methylation-associated SNPs from a regulatory perspective, we used DeepSEA ^42^, a machine learning algorithm that predicts SNP functionality from the sequence context based on sequencing data for different regulatory elements in different cell lines using ENCODE data ^66^. We identified the SNPs with putatively functional consequences on regulatory marks by DeepSEA and compared putative regulatory effects of G, G+E and GxE hits. Relative to the imputed non-DeepSEA SNPs contained in our dataset, these predicted functional DeepSEA SNPs were enriched for TSS and enhancer regions and depleted for quiescent regions, supporting their relevance in regulatory processes (see Figure 4). Compared to DeepSEA SNPs overall, DeepSEA SNPs within the 3 different best models also showed distinct enrichment or depletion patterns. Likely functional GxE SNPs showed, similar to GxE VMRs, relative depletion in TSS regions while G+E SNPs showed enrichment in genic enhancers. Overall, both the VMRs as well as the associated functional SNPs appear to be in distinct regulatory regions, depending on their best model. In addition, GxE functional SNP and tagCpGs were located farther apart than SNP/tagCpG pairs within G or G+E models (see Figure S2B), supporting a more long-range type of regulation in GxE interactions on molecular traits as compared to all genes; a similar relationship has been reported previously for GxE with regard to gene expression in *C. elegans* ^89; 90^.

In addition, eQTMs ^68^ were also significantly enriched for VMRs selected in these study as compared to randomly selected CpGs, supporting that differential methylation of these sites, driven by genetic and environmental factors can correlate with gene expression. SNPs associated with differences in gene expression but also DNA methylation have consistently been shown to be enriched among SNPs associated with common disorders in GWAS ^24; 27; 29;50^. This is in line with the fact that functionally relevant SNPs chosen by DeepSEA, selected for their predicted effects on transcriptional and epigenetic regulation, are themselves significantly enriched in GWAS for all tested common disorders. On top of this baseline enrichment of all DeepSEA SNPs, the genetic variants that were within G, GxE or G+E models predicting variable DNA methylation were even further enriched in GWAS association results, but again with some differences in the pattern of enrichment. GxE DeepSEA SNPs were most enriched in GWAS association for three psychiatric disorders, i.e. autism spectrum disorders, bipolar disorder and major depressive disorder, while SNPs within G+E models were most enriched in GWAS for inflammatory bowel disease. The fact that the enrichment was observed for not only G and G+E SNPs, with strong main genetic effects, but also for GxE SNPs, with smaller to sometimes no main genetic effect on DNA methylation underscores the importance of also including SNPs within GxE models in the functional annotation of GWAS. A detailed catalogue of meQTLs that are responsive to prenatal environments could help support a better pathophysiological understanding of diseases for which risk is shaped by a combination of prenatal environment and genetic factors.

Finally, we want to note the limitations of this study. First, we restricted our analyses to specific DNA methylation array contents that are inherently biased as compared to genome-wide bisulfite sequencing, for example. In addition, we restricted our analysis to VMRs. Ong and Holbrooke ^54^ showed that this approach increases statistical power. Furthermore, VMRs are enriched for enhancers and transcription factor binding sites, overlap with GWAS hits ^67^ and are associated with gene-expression of nearby genes at these sites ^91^. This is in line with our observation that identified VMRs presented with intermediate methylation levels which have been shown to be enriched in regions of regulatory function, like enhancers, exons and DNase I hypersensitivity sites ^92^. Hence, the effects of genotypes on DNA methylation levels in VMRs might be higher as compared to less variable CpG-sites. In addition, genotypes are measured with much less error as compared to environmental factors which may also reduce the overall explained variance in large cohorts.

Second, it has been reported that different cell types display different patterns of DNA methylation ^67^. Therefore, the most variable CpG-sites may also include those that reflect differences in cord blood cell type proportions. To address this issue, all analyses were corrected for estimated cell proportions to the best of our current availability, so that differences in cell type proportion likely do not account for all of the observed effects. However, only replication in specific cell types will be able to truly assess the proportion of VMRs influenced by this.

Third, we used the AIC as main criterion for model fit ^64^ which is equivalent to a penalized likelihood-function. There are a variety of other model selection criteria ^93^ and choosing between these is an ongoing debate which also depends on the underlying research question. We decided to use the AIC as one of our main aims was to compare our results with the study of Teh et al. ^34^ in which this criterion was applied and as this method maybe more powerful for detecting GxE than for example model selection criteria based on lowest p-values.

Finally, our analyses are restricted to DNA methylation in neonatal blood and to pregnancy environments. Whether similar conclusions can be drawn for methylation levels assessed at a later developmental stage needs to be investigated.

## Conclusion

We tested whether genotype, a combination of different prenatal environmental factors and the additive or the multiplicative interactive effects of both mainly influence VMRs in the newborn’s epigenome. Our results show that G in combination with E are the best predictors of variance in DNA methylation. This highlights the importance of including both individual genetic differences as well as environmental phenotypes into epigenetic studies and also the importance of improving our ability to identify environmental associations. Our data also support the disease relevance of variants predicting DNA methylation together with the environment beyond main meQTL effects, and the view that there are functional differences of additive and interactive effects of genes and environment on DNA methylation. Improved understanding of these functional differences may also yield novel insights into pathophysiological mechanisms of common non-communicable diseases, as risk for all of these disorders is driven by both genetic and environmental factors.

## Methods

### Availability of data and materials

The datasets analyzed during the current study are not publicly available. However,an interested researcher can obtain a de-identified dataset after approval from the PREDO Study Board. Data requests may be subject to further review by the national register authority and by the ethical committees. Any requests for data use should be addressed to the PREDO Study Board or individual researchers.

### The PREDO cohort

The Prediction and Prevention of Preeclampsia and Intrauterine Growth Restriction (PREDO) Study is a longitudinal multicenter pregnancy cohort study of Finnish women and their singleton children born alive between 2006-2010 ^35^. We recruited 1,079 pregnant women, of whom 969 had one or more and 110 had none of the known clinical risk factors for preeclampsia and intrauterine growth restriction. The recruitment took place when these women attended the first ultrasound screening at 12+0-13+6 weeks+days of gestation in one of the ten hospital maternity clinics participating in the study. The cohort profile ^35^ contains details of the study design and inclusion criteria.

### Ethics

The study protocol was approved by the Ethical Committees of the Helsinki and Uusimaa Hospital District and by the participating hospitals. A written informed consent was obtained from all women.

### Maternal characteristics

We tested 10 different maternal environments:

### Depressive symptoms

Starting from 12+0-13+6 gestational weeks+days pregnant women filled in the 20 item Center for Epidemiological Studies Depression Scale (CES-D) ^43^ for depressive symptoms in the past 7 days. They filled in the CES-D scale biweekly until 38+0-39+6 weeks+days of gestation or delivery. We used the mean-value across all the CES-D measurements.

### Symptoms of anxiety

At 12+0-13+6 weeks+days of gestation, women filled in the 20 item Spielberger’s State Trait Anxiety Inventory (STAI) ^44^ for anxiety symptoms in the past 7 days. They filled in the STAI scale biweekly until 38+0-39+6 weeks+days of gestation or delivery. We used the mean-value across all these measurements.

### Betamethasone

Antenatal betamethasone treatment (yes/no) was derived from the hospital records and the Finnish Medical Birth Register (MBR).

### Delivery method

Mode of delivery (vaginal delivery vs. caesarean section) was derived from patient records and MBR.

### Parity

Parity (number of previous pregnancies leading to childbirth) at the start of present pregnancy was derived from the hospital records and the MBR.

### Maternal age

Maternal age at delivery (years) was derived from the hospital records and the MBR.

### Pre-pregnancy BMI

Maternal pre-pregnancy BMI (kg/m^2^), calculated from measurements weight and height verified at the first antenatal clinic visit at 8+4 (SD 1+3) gestational week was derived from the hospital records and the MBR.

### Hypertension

Hypertension was defined as any hypertensive disorder including gestational hypertension, chronic hypertension and preeclampsia against normotension. Gestational hypertension was defined as systolic/diastolic blood pressure ≥140/90 mm Hg on ≥ 2 occasions at least 4 h apart in a woman who was normotensive before 20^th^ week of gestation. Preeclampsia was defined as systolic/diastolic blood pressure ≥140/90 mm Hg on ≥2 occasions at least 4 h apart after 20^th^ week of gestation and proteinuria ≥300 mg/24 h. Chronic hypertension was defined as systolic/diastolic blood pressure ≥140/90 mm Hg on ≥2 occasions at least 4 h apart before 20^th^ gestational week or medication for hypertension before 20 weeks of gestation.

### Gestational diabetes and oral glucose tolerance test (OGTT)

Gestational diabetes was defined as fasting, 1h or 2h plasma glucose during a 75g oral glucose tolerance test ≥5.1, ≥10.0 and/or ≥8.5 mmol/L, respectively, that emerged or was first identified during pregnancy. We took the area under the curve from the three measurements as a single measure for the OGTT itself.

### Genotyping and Imputation

Genotyping was performed on Illumina Human Omni Express Exome Arrays containing 964,193 SNPs. Only markers with a call rate of at least 98%, a minor allele frequency of at least 1% and a p-value for deviation from Hardy-Weinberg-Equilibrium > 1.0 × 10^−06^ were kept in the analysis. After QC, 587,290 SNPs were available.

In total, 996 cord blood samples were genotyped. Samples with a call rate below 98% (n=11) were removed.

Any pair of samples with IBD estimates > 0.125 was checked for relatedness. As we corrected for admixture in our analyses, these samples were kept except for one pair which could not be resolved. From this pair we excluded one sample from further analysis.

Individuals showing discrepancies between phenotypic and genotypic sex (n=1) were removed. We also checked for heterozygosity outliers but found none. 983 participants were available in the final dataset.

Before imputation, AT and CG SNPs were removed. Imputation was performed using shapeit2 (https://mathgen.stats.ox.ac.uk./genetics_software/shapeit/shapeit.html) and impute2 (https://mathgen.stats.ox.ac.uk/impute/impute_v2.html). Chromosomal and base pair positions were updated to the 1000 Genomes Phase 3 reference set, allele strands were flipped where necessary.

After imputation, we reran quality control, filtering out SNPs with an info score < 0.8, a minor allele frequency below 1% and a deviation from HWE with a p-value < 1.0 × 10^−06^. This resulted in a dataset of 9,402,991 SNPs. After conversion into best guessed genotypes using a probability threshold of 90%, we performed another round of QC (using SNP-call rate of least 98%, a MAF of at least 1% and a p-value threshold for HWE of 1.0 × 10^−06^), after which 7,314,737 SNPs remained for the analysis.

For the evaluation of which model best explained the methylation sites, we pruned the dataset using a threshold of r^2^ of 0.2 and a window-size of 50 SNPs with an overlap of 5 SNPs. The final, pruned dataset contained 788,156 SNPs. 36,241 of these variants were DeepSea variants (see Methods below).

### Methylation

Cord blood samples were run on Illumina 450k Methylation arrays. The quality control pipeline was set up using the R-package *minfi* ^45^ (https://www.r-project.org). Three samples were excluded as they were outliers in the median intensities. Furthermore, 20 samples showed discordance between phenotypic sex and estimated sex and were excluded. Nine samples were contaminated with maternal DNA according to the method suggested by Morin et al. ^46^ and were also removed.

Methylation beta-values were normalized using the *funnorm* function ^47^. After normalization, two batches, i.e., slide and well, were significantly associated and were removed iteratively using the *Combat* function ^48^ in the *sva* package ^49^.

We excluded any probes on chromosome X or Y, probes containing SNPs and cross-hybridizing probes according to Chen et al. ^50^ and Price et al ^51^. Furthermore, any CpGs with a detection p-value > 0.01 in at least 25% of the samples were excluded.

The final dataset contained 428,619 CpGs and 822 participants. For 817 of these, also genotypes were available.

An additional 161 cord blood samples were run on Illumina EPIC Methylation arrays. Three samples were excluded as they were outliers in the median intensities. Three samples showed discordance between phenotypic sex and estimated sex and were excluded. Three samples were contaminated with maternal DNA and were also removed ^46^.

Methylation beta-values were normalized using the *funnorm* function ^47^ in the R*–*package *minfi ^45^*. Three samples showed density artefacts after normalization and were removed from further analysis. We excluded any probes on chromosome X or Y, probes containing SNPs and cross-hybridizing probes according to Chen et al. ^50^, Price et al. ^51^ and McCartney et al. ^52^. Furthermore, any CpGs with a detection p-value > 0.01 in at least 25% of the samples were excluded. The final dataset contains 812,987 CpGs and 149 samples. After normalization no significant batches were identified. For 146 of these samples, genotypic data was also available.

Cord blood cell counts were estimated for seven cell types (nucleated red blood cells, granulocytes, monocytes, natural killer cells, B cells, CD4(+)T cells, and CD8(+)T cells) using the method of Bakulski et al. ^53^ which is incorporated in the R-package *minfi ^45^*.

### Identification of VMRs (variable methylated regions)

The VMR approach was described by Ong and Holbrook ^54^. We chose all 42,862 CpGs with a MAD score greater than the 90^th^ percentile. A VMR region was defined as at least two spatially contiguous probes which were at most 1kb apart of each other. This resulted in 3,982 VMRs in the 450K samples and in 8,547 VMRs in the EPIC sample. The CpG with the highest MAD scores was chosen as representative of the whole VMR in the statistical analysis.

### Drakenstein cohort

Details on this cohort and the assessed phenotypes can be found in ^39; 40^. The birth cohort design recruits pregnant women attending one of two primary health care clinics in the Drakenstein sub-district of the Cape Winelands, Western Cape, South Africa – Mbekweni (serving a black African population) and TC Newman (serving a mixed ancestry population). Consenting mothers were enrolled during pregnancy, and mother–child dyads are followed longitudinally until children reach at least 5 years of age. Mothers are asked to request that the father of the index pregnancy attend a single antenatal study visit where possible. Follow-up visits for mother–child dyads take place at the two primary health care clinics and at Paarl Hospital.

Pregnant women were eligible to participate if they were 18 years or older, were accessing one of the two primary health care clinics for antenatal care, had no intention to move out of the district within the following year, and provided signed written informed consent. Participants were enrolled between 20 and 28 weeks’ gestation, upon presenting for antenatal care visit. In addition, consenting fathers of the index pregnancy when available were enrolled in the study and attended a single antenatal study visit.

### Ethics

The study was approved by the Faculty of Health Sciences, Human Research Ethics Committee, University of Cape Town (401/2009), by Stellenbosch University (N12/02/0002), and by the Western Cape Provincial Health Research committee (2011RP45). All participants provided written informed consent.

### Maternal characteristics

After providing consent, participants were asked to complete a battery of self-report and clinician-administered measures at a number of antenatal and postnatal study visits. All assessed phenotypes are described in detail in ^39^. Here, we give a short outline on the phenotypes which were used in our analysis. Maternal parity was obtained from the antenatal record; maternal age was from the date of birth as recorded on the mothers’ national identity document. The mode of delivery was ascertained by direct observation of the birth by a member of the study team as all births occurred at Paarl hospital. The SRQ-20 ^55^ is a WHO-endorsed measure of psychological distress consisting of 20 items which assess non-psychotic symptoms, including symptoms of depressive and anxiety disorders. Each item is scored according to whether the participant responds in the affirmative (scored as 1) or negative (scored as 0) to the presence of a symptom. Individual items are summed to generate a total score. The Beck Depression Inventory (BDI-II) is a widely-used and reliable measure of depressive symptoms ^56^. The BDI-II comprises 21 items, each of which assesses the severity of a symptom of major depression. Each item is assessed on a severity scale ranging from 0 (absence of symptoms) to 3 (severe, often with functional impairment). A total score is then obtained by summing individual item responses, with a higher score indicative of more severe depressive symptoms.

Smoking was assessed using The Alcohol, Smoking and Substance Involvement Screening Test (ASSIST) ^57^, a tool that was developed by the WHO to detect and manage substance use among people attending primary health care services. The tool assesses substance use and substance-related risk across 10 categories (tobacco, alcohol, cannabis, cocaine, amphetamine-type stimulants, inhalants, sedatives/sleeping pills, hallucinogens, opioids, and other substances), as well as enquiring about a history of intravenous drug use. Total scores are obtained for each substance by summing individual item responses, with a higher score indicative of greater risk for substance-related health problems.

Hypertension was assessed by blood pressure measured antenatally.

### Genotyping and Imputation

Genotyping in DCHS was performed using the Illumina PsychArray for those samples with 450k data, or the Illumina GSA for those samples with EPIC DNA methylation data (Illumina, San Diego, USA). For both array types, QC and imputation was the same; first, raw data was imported into Genome Studio and exported into R for QC. SNPs were filtered out if they had a tenth percentile GC score below 0.2 or an average GC score below 0.1, for a total of 140 SNPs removed. Phasing was performed using shapeit, and imputation was performed using impute2 with 1000 Genomes Phase 1 reference data. After imputation, we used qctool to filter out SNPs with an info score <0.8 or out of Hardy-Weinberg equilibrium.

As after imputation, only 5,286 DeepSEA variants were available for those samples genotyped on the PsychArray and only 4,049 for those samples genotyped on the GSAchip, we performed LD-pruning based on a threshold of r^2^ of 0.2 and a window-size of 50 SNPs with an overlap of 5 SNPs. This resulted in 162,292 SNPs (PsychArray) and 176,553 SNPs (GSAchip).

### Methylation

Details for DNA methylation measurements and quality control have been previously published ^46^. Briefly, we performed basic quality control on data generated by either the 450k or EPIC arrays using Illumina’s Genome Studio software for background subtraction and colour correction. Data was filtered to remove CpGs with high detection p values, those on the X or Y chromosome, or with previously identified poor performance. 450k data was normalized using SWAN and EPIC data using BMIQ, and both used ComBat to correct for chip (both), and row (450k only). The final analysis was performed with 107 samples with methylation levels from the 450k array and 151 with methylation levels assessed on the EPIC array and available genotypes.

### VMRs

We identified 6,072 VMRs in DCHS I and 10,005 VMRs in DCHS II.

### The UCI cohort

Mothers and children were part of an ongoing, longitudinal study, conducted at the University of California, Irvine (UCI), for which mothers were recruited during the first trimester of pregnancy ^36-38^. All women had singleton, intrauterine pregnancies. Women were not eligible for study participation if they met the following criteria: corticosteroids, or illicit drugs during pregnancy (verified by urinary cotinine and drug toxicology). Exclusion criteria for the newborn were preterm birth (i.e., less than 34 weeks of gestational age at birth), as well as any congenital, genetic, or neurologic disorders at birth.

### Ethics

The UCI institutional review board approved all study procedures and all participants provided written informed consent.

### Maternal Characteristics

Maternal sociodemographic characteristics (age, parity) were obtained via a standardized structured interview at the first pregnancy visit. Maternal pre-pregnancy BMI (weight kg/height m^2^) was computed based on pre-pregnancy weight abstracted from the medical record, and maternal height was measured at the research laboratory during the first pregnancy visit. Obstetric risk conditions during pregnancy, including presence of gestational diabetes and hypertension, and delivery mode were abstracted from the medical record. At each pregnancy visit the Center for Epidemiological Studies Depression Scale ^43^ and the State scale from the State–Trait Anxiety Inventory ^44^ were administered. For individuals with <3 missing items on any scale at any time point, the mean responses for that scale were calculated and then multiplied by the total number of items in the respective scale, to generate total scale scores that are comparable to those generated from participants without any missing data. We used the average depression and anxiety score throughout pregnancy in the calculations. Maternal smoking during pregnancy was determined by maternal self-report and verified by measurement of urinary cotinine concentration. Urinary cotinine was assayed in maternal samples collected at each trimester using the Nicotine/COT(Cotinine)/Tobacco Drug Test Urine Cassette (http://www.meditests.com/nicuintescas.html), which involves transferring 4 drops of room temperature urine into the well of the cassette, and employs a cutoff for COT presence of 200ng/ml. Endorsement of smoking or detection of urinary COT in any trimester was coded as 1, and absence of evidence for smoking in any trimester coded as 0.

### Genotyping

Genomic DNA was extracted from heel prick blood samples and used for all genomic analysis. Genotyping was performed on Illumina Human Omni Express (24 v1.1) Arrays containing 713,014 SNPs. All samples had a high call rate (above 97%). SNPs with a minor allele frequency >5% and a p-value for deviation from Hardy-Weinberg-Equilibrium > 1.0 × 10^−25^ were retained for analysis. After QC, 602,807 SNPs were available.

### Imputation

Before imputation, chromosomal and base pair positions were updated to the Haplotype Reference Consortium (r1.1) reference set, allele strands were flipped where necessary. Phasing was performed using EAGLE2 (https://data.broadinstitute.org/alkesgroup/Eagle/) and imputation was performed using PBWT (https://github.com/VertebrateResequencing/pbwt). Imputed SNPs with an info score < 0.8, duplicates and ambiguous SNPs were removed resulting in 21,341,980 SNPs. Of these, 19,530 were DeepSEA variants.

### DNA Methylation

DNAm analysis using the Infinium Illumina MethylationEPIC BeadChip (Illumina, Inc., San Diego, CA) was performed according to the manufacturer’s guidelines in using genomic DNA derived from neonatal heel prick samples. Quality Control carried out in *minfi* ^45^. No outliers were detected in the median intensities of methylated and unmethylated channels. All samples had a high call rate of at least 95% and their predicted sex was the same as the phenotypic sex. We removed CpGs with a high detection value (p<0.0001), probes missing >3 beads in >5% of the cohort, in addition to non-specific/cross-hybridizing and SNP probes ^51; 52^. Methylation beta-values were normalized using functional normalization (*funnorm*) ^47^.

We also iteratively adjusted the data for relevant technical factors, i.e. array row, experimental batch and sample plate, using *Combat* ^48^. The final dataset contained 768,910 CpGs. Neonatal blood cell counts were estimated for seven cell types: nucleated red blood cells, granulocytes, monocytes, natural killer cells, B cells, CD4(+)T cells, and CD8(+)T cells ^53^.

The final dataset contained 121 samples with available genotypes and methylation values. *VMRs*

Applying the same procedure as for PREDO I and PREDO II, we identified 9,525 VMRs in the ICU cohort.

### MoBa cohort

Participants represent two subsets of mother-offspring pairs from the national Norwegian Mother and Child Cohort Study (MoBa) ^58^. MoBa is a prospective population-based pregnancy cohort study conducted by the Norwegian Institute of Public Health. The years of birth for MoBa participants ranged from 1999-2009. MoBa mothers provided written informed consent. Each subset is referred to here as MoBa1 and MoBa2. MoBa1 is a subset of a larger study within MoBa that included a cohort random sample and cases of asthma at age three years ^59^. We previously reported an association between maternal smoking during pregnancy and differential DNA methylation in MoBa1 newborns ^60^. We subsequently measured DNA methylation in additional newborns (MoBa2) in the same laboratory (Illumina, San Diego, CA) ^11^. MoBa2 included cohort random sample plus cases of asthma at age seven years and non-asthmatic controls. Years of birth were 2002-2004 for children in MoBa1, 2000-2005 for MoBa2.

### Ethics

The establishment and data collection in MoBa obtained a license from the Norwegian Data Inspectorate and approval from The Regional Committee for Medical Research Ethics. Both studies were approved by the Regional Committee for Ethics in Medical Research, Norway. In addition, MoBa1 and MoBa2 were approved by the Institutional Review Board of the National Institute of Environmental Health Sciences, USA.

### Maternal characteristics

To replicate specific GxE and G+E from PREDO I, we focused on those characteristics which were available in both cohorts: maternal age, pre-pregnancy BMI and hypertension.

Within MoBa, the questionnaires at weeks 17 and 30 include general background information as well as details on previous and present health problems and exposures. The birth record from the Medical Birth Registry of Norway ^61^which includes maternal health during pregnancy as well as procedures around birth and pregnancy outcomes, is integrated in the MoBa database.

### Genotyping and Imputation

DNA was extracted from the MoBa biobank and genotyped on the Illumina HumanExomeCore platform. The genotypes were called with GenomeStudio software. Phasing and imputation were done using shapeit2 (https://mathgen.stats.ox.ac.uk./genetics_software/shapeit/shapeit.html) and impute2 (https://mathgen.stats.ox.ac.uk/impute/impute_v2.html) with the thousand genomes phase 3 reference panel for the European population. Variates with a imputation score of less than 0.8 were filtered out.

### Methylation

Details of the DNA methylation measurements and quality control for the MoBa1 participants were previously described ^41^ and the same protocol was implemented for the MoBa2 participants. Briefly, at birth, umbilical cord blood samples were collected and frozen at birth at −80**°**C. All biological material was obtained from the Biobank of the MoBa study ^41^.

Bisulfite conversion was performed using the EZ-96 DNA Methylation kit (Zymo Research Corporation, Irvine, CA) and DNA methylation was measured at 485,577 CpGs in cord blood using Illumina’s Infinium HumanMethylation450 BeadChip ^62^. Raw intensity (.idat) files were handled in R using the *minfi* package to calculate the methylation level at each CpG as the beta-value (β=intensity of the methylated allele (M)/(intensity of the unmethylated allele (U) + intensity of the methylated allele (M) + 100)) and the data was exported for quality control and processing. Control probes (N=65) and probes on X (N=11 230) and Y (N=416) chromosomes were excluded in both datasets. Remaining CpGs missing > 10% of methylation data were also removed (N=20 in MoBa1, none in MoBa2). Samples indicated by Illumina to have failed or have an average detection p value across all probes < 0.05 (N=49 MoBa1, N=35 MoBa2) and samples with gender mismatch (N=13 MoBa1, N=8 MoBa2) were also removed. For MoBa1 and MoBa2, we accounted for the two different probe designs by applying the intra-array normalization strategy Beta Mixture Quantile dilation (BMIQ) ^63^.

The Empirical Bayes method via *ComBat* was applied separately in MoBa1 and MoBa2 for batch correction using the *sva* package in R ^49^. After quality control exclusions, the sample sizes were 1,068 for MoBa1 and 685 for MoBa2.

After QC, the total number of samples was 1,732, with 1,592 overlapping with the methylation samples.

### Regression analysis

Regression analysis was conducted using the lm function in R 3.3.1 (https://www.r-project.org). We included the child’s sex, seven estimated cell counts as well as the first two (PREDO I and PREDO II), first three (UCI) and first five (DCHS I and II) principal components of the MDS analysis on the genotypes in the model, the corresponding plot of the first ten MDS-components in PREDO is depicted in Figure S4. SNP genotypes were recoded into a count of 0, 1 or 2 representing the number of minor allele copies. For each VMR site, we tested SNPs located in a 1MB window up- and downstream of the specific site. In PREDO and UCI, we restricted the analysis to DeepSEA variants while we used the pruned SNP-set in DCHS.

For each VMR we tested four models:

1. Methylation ~ covariates + environment
2. Methylation ~ covariates + SNP
3. Methylation ~ covariates + SNP + environment
4. Methylation ~ covariates + SNP + environment + SNP x environment

For each model, the AIC, Akaike’s information criterion ^64^ was calculated and the model with the lowest AIC was chosen as the best model.

### Enrichment analyses

With regard to enrichment for VMRs, CpG-site within VMRs were compared to all other CpG-sites on the 450K array located in non-VMR-regions. With regard to enrichment for VMRs best explained by G, G+E or GxE, tagCpGs best explained by the specific model were compared to tagCpGs best explained by any of the other models. For enrichment tests for DeepSEA SNPs, non-DeepSEA SNPs present in our dataset were used as comparison group. Enrichment tests were performed based on a hyper-geometric test, i.e. a Fisher-test. The significance levels was set at p<0.05.

With regard to enrichment for GWAS hits, DeepSEA variants were matched to GWAs variants based on chromosome and position (hg19). To check for enrichment for nominal significant GWAS hits, the full summary statistics were derived from the respective publication.

Histone ChiP-seq peaks from Roadmap Epigenomics project for blood and embryonic stem cells were downloaded from http://egg2.wustl.edu/roadmap/data/byFileType/peaks/consolidated/broadPeak/. The pre-processed consolidated broad peaks from the uniform processing pipeline of the Roadmap project were used.

### Genomic annotation mapping

CpG sites were mapped to the genome location according to Illumina’s annotation using the R-package *minfi*.

### DeepSEA Analysis

Pretrained DeepSEA model was downloaded from: http://deepsea.princeton.edu/media/code/deepsea.v0.94.tar.gz and variant files in VCF format are used for producing e-values. VCF files were first split into smaller files each containing one million variants and the model was run using the command line on a server with a NVIDIA Titan X GPU card.

We reran our models using only DeepSEA variants which had been identified by the algorithm of Zhou and Troyanskaya ^42^. This method predicts functionality of a SNP based on the DNA-sequence. We included all 212,210 variants with a functional significance e-value below 5 × 10^−05^. The e-values represent the significance of the regulatory impact of given variants compared to one million random variants.

### Random-effects meta-analysis

GxE and G+E result for PREDO and for MoBa were meta-analysed using a random-effects model in the R-package rmeta. Replication was defined as DeepSEA-tagCpG-environment combinations showing the same effect direction in both cohorts, presenting with smaller p-values as for PREDO alone and with a FDR-corrected p-value (across all combinations tested in the meta-analysis) below 0.05.

### Consortia

<u>Major Depressive Disorder Working Group of the Psychiatric Genomics Consortium</u>: a full list of members as well as their affiliations is given in the Supplemental Information.

## Acknowledgements

We want to thank Susanne Sauer and Maik Ködel for their technical assistance. We thank all mothers who took part in the on-going PREDO study. We are grateful to all the families in Norway who participate in the on-going MoBa cohort study. We thank the Drakenstein Child Health Study staff, and the clinical and administrative staff of the Western Cape Government Department of Health at Paarl Hospital and at the clinics for support of the Study. We also thank our collaborators and students. Finally, we thank all mothers and children enrolled in the Drakenstein Child Health Study. We thank the research participants and employees of 23andMe, Inc. for their contribution to this study.

## Author’s contributions

DC and EBB conceived the analyses.

JL, MLP, EH, EK, HL, PMV, RRM and KR conceptualized and planned the PREDO study and collected the data.

CMP, WN, SH and SJL conceptualized and planned the MoBa study and collected the data. CB, SE, PWD, KJOD conceptualized and planned the UCI study and collected the data.

DTSL and JLM performed the DNA methylation and genotyping arrays for the UCI and DCH studies.

DJS, NK, HJZ designed and undertook the DCHS; MJM, MSK, KCK were involved in testing and analysis of epigenetic data; SD was involved in testing and analysis of genetic data.

DC, GE, CMP and MJJ ran the statistical analysis. NSM, IK and FJT co-supervised statistical analysis.

DC and EBB wrote the manuscript with contributions from GE, SJL, CMP, KR, JL. DC, JL, KR, and EB interpreted the results.

All authors contributed to and approved the final version of the manuscript.

## Availability of data and materials

The datasets analyzed during the current study are not publicly available. However, an interested researcher can obtain a de-identified dataset after approval from the PREDO Study Board. Data requests may be subject to further review by the national register authority and by the ethical committees. Any requests for data use should be addressed to the PREDO Study Board or individual researchers.

## Bibliography

1. Roseboom, T., de Rooij, S., and Painter, R. (2006). The Dutch famine and its long-term consequences for adult health. Early Hum Dev 82, 485–491.

2. Barker, D.J., Osmond, C., Forsen, T.J., Kajantie, E., and Eriksson, J.G. (2005). Trajectories of growth among children who have coronary events as adults. N Engl J Med 353, 1802–1809.

3. Hovi, P., Andersson, S., Eriksson, J.G., Jarvenpaa, A.L., Strang-Karlsson, S., Makitie, O., and Kajantie, E. (2007). Glucose regulation in young adults with very low birth weight. N Engl J Med 356, 2053–2063.

4. Hillier, T.A., Pedula, K.L., Schmidt, M.M., Mullen, J.A., Charles, M.A., and Pettitt, D.J. (2007). Childhood obesity and metabolic imprinting: the ongoing effects of maternal hyperglycemia. Diabetes Care 30, 2287–2292.

5. Dancause, K.N., Laplante, D.P., Hart, K.J., O’Hara, M.W., Elgbeili, G., Brunet, A., and King, S. (2015). Prenatal stress due to a natural disaster predicts adiposity in childhood: the Iowa Flood Study. J Obes 2015, 570541.

6. Lahti, M., Savolainen, K., Tuovinen, S., Pesonen, A.K., Lahti, J., Heinonen, K., Hamalainen, E., Laivuori, H., Villa, P.M., Reynolds, R.M., et al. (2017). Maternal Depressive Symptoms During and After Pregnancy and Psychiatric Problems in Children. J Am Acad Child Adolesc Psychiatry 56, 30–39 e37.

7. Bronson, S.L., and Bale, T.L. (2016). The Placenta as a Mediator of Stress Effects on Neurodevelopmental Reprogramming. Neuropsychopharmacology 41, 207–218.

8. Schwarze, C.E., Mobascher, A., Pallasch, B., Hoppe, G., Kurz, M., Hellhammer, D.H., and Lieb, K. (2013). Prenatal adversity: a risk factor in borderline personality disorder? Psychol Med 43, 1279–1291.

9. Entringer, S., Buss, C., and Wadhwa, P.D. (2015). Prenatal stress, development, health and disease risk: A psychobiological perspective-2015 Curt Richter Award Paper. Psychoneuroendocrinology 62, 366–375.

10. Gutierrez-Arcelus, M., Lappalainen, T., Montgomery, S.B., Buil, A., Ongen, H., Yurovsky, A., Bryois, J., Giger, T., Romano, L., Planchon, A., et al. (2013). Passive and active DNA methylation and the interplay with genetic variation in gene regulation. Elife 2, e00523.

11. Joubert, B.R., Felix, J.F., Yousefi, P., Bakulski, K.M., Just, A.C., Breton, C., Reese, S.E., Markunas, C.A., Richmond, R.C., Xu, C.J., et al. (2016). DNA Methylation in Newborns and Maternal Smoking in Pregnancy: Genome-wide Consortium Meta-analysis. Am J Hum Genet 98, 680–696.

12. Joubert, B.R., den Dekker, H.T., Felix, J.F., Bohlin, J., Ligthart, S., Beckett, E., Tiemeier, H., van Meurs, J.B., Uitterlinden, A.G., Hofman, A., et al. (2016). Maternal plasma folate impacts differential DNA methylation in an epigenome-wide meta-analysis of newborns. Nat Commun 7, 10577.

13. Sharp, G.C., Salas, L.A., Monnereau, C., Allard, C., Yousefi, P., Everson, T.M., Bohlin, J., Xu, Z., Huang, R.C., Reese, S.E., et al. (2017). Maternal BMI at the start of pregnancy and offspring epigenome-wide DNA methylation: findings from the pregnancy and childhood epigenetics (PACE) consortium. Hum Mol Genet 26, 4067–4085.

14. Girchenko, P., Lahti, J., Czamara, D., Knight, A.K., Jones, M.J., Suarez, A., Hamalainen, E., Kajantie, E., Laivuori, H., Villa, P.M., et al. (2017). Associations between maternal risk factors of adverse pregnancy and birth outcomes and the offspring epigenetic clock of gestational age at birth. Clin Epigenetics 9, 49.

15. Rijlaarsdam, J., Pappa, I., Walton, E., Bakermans-Kranenburg, M.J., Mileva-Seitz, V.R., Rippe, R.C., Roza, S.J., Jaddoe, V.W., Verhulst, F.C., Felix, J.F., et al. (2016). An epigenome-wide association meta-analysis of prenatal maternal stress in neonates: A model approach for replication. Epigenetics 11, 140–149.

16. Martin, E.M., and Fry, R.C. (2018). Environmental Influences on the Epigenome: Exposure-Associated DNA Methylation in Human Populations. Annu Rev Public Health 39, 309–333.

17. El Hajj, N., Haertle, L., Dittrich, M., Denk, S., Lehnen, H., Hahn, T., Schorsch, M., and Haaf, T. (2017). DNA methylation signatures in cord blood of ICSI children. Hum Reprod 32, 1761–1769.

18. Sosnowski, D.W., Booth, C., York, T.P., Amstadter, A.B., and Kliewer, W. (2018). Maternal prenatal stress and infant DNA methylation: A systematic review. Dev Psychobiol 60, 127–139.

19. Bauer, T., Trump, S., Ishaque, N., Thurmann, L., Gu, L., Bauer, M., Bieg, M., Gu, Z., Weichenhan, D., Mallm, J.P., et al. (2016). Environment-induced epigenetic reprogramming in genomic regulatory elements in smoking mothers and their children. Mol Syst Biol 12, 861.

20. Sharp, G.C., Lawlor, D.A., Richmond, R.C., Fraser, A., Simpkin, A., Suderman, M., Shihab, H.A., Lyttleton, O., McArdle, W., Ring, S.M., et al. (2015). Maternal pre-pregnancy BMI and gestational weight gain, offspring DNA methylation and later offspring adiposity: findings from the Avon Longitudinal Study of Parents and Children. Int J Epidemiol 44, 1288–1304.

21. Lin, X., Lim, I.Y., Wu, Y., Teh, A.L., Chen, L., Aris, I.M., Soh, S.E., Tint, M.T., MacIsaac, J.L., Morin, A.M., et al. (2017). Developmental pathways to adiposity begin before birth and are influenced by genotype, prenatal environment and epigenome. BMC Med 15, 50.

22. Cecil, C.A., Walton, E., Smith, R.G., Viding, E., McCrory, E.J., Relton, C.L., Suderman, M., Pingault, J.B., McArdle, W., Gaunt, T.R., et al. (2016). DNA methylation and substance-use risk: a prospective, genome-wide study spanning gestation to adolescence. Transl Psychiatry 6, e976.

23. Gibbs, J.R., van der Brug, M.P., Hernandez, D.G., Traynor, B.J., Nalls, M.A., Lai, S.L., Arepalli, S., Dillman, A., Rafferty, I.P., Troncoso, J., et al. (2010). Abundant quantitative trait loci exist for DNA methylation and gene expression in human brain. PLoS Genet 6, e1000952.

24. Gaunt, T.R., Shihab, H.A., Hemani, G., Min, J.L., Woodward, G., Lyttleton, O., Zheng, J., Duggirala, A., McArdle, W.L., Ho, K., et al. (2016). Systematic identification of genetic influences on methylation across the human life course. Genome Biol 17, 61.

25. McClay, J.L., Shabalin, A.A., Dozmorov, M.G., Adkins, D.E., Kumar, G., Nerella, S., Clark, S.L., Bergen, S.E., Swedish Schizophrenia, C., Hultman, C.M., et al. (2015). High density methylation QTL analysis in human blood via next-generation sequencing of the methylated genomic DNA fraction. Genome Biol 16, 291.

26. Chen, L., Ge, B., Casale, F.P., Vasquez, L., Kwan, T., Garrido-Martin, D., Watt, S., Yan, Y., Kundu, K., Ecker, S., et al. (2016). Genetic Drivers of Epigenetic and Transcriptional Variation in Human Immune Cells. Cell 167, 1398–1414 e1324.

27. Hannon, E., Weedon, M., Bray, N., O’Donovan, M., and Mill, J. (2017). Pleiotropic Effects of Trait-Associated Genetic Variation on DNA Methylation: Utility for Refining GWAS Loci. Am J Hum Genet 100, 954–959.

28. Pierce, B.L., Tong, L., Argos, M., Demanelis, K., Jasmine, F., Rakibuz-Zaman, M., Sarwar, G., Islam, M.T., Shahriar, H., Islam, T., et al. (2018). Co-occurring expression and methylation QTLs allow detection of common causal variants and shared biological mechanisms. Nat Commun 9, 804.

29. Cheung, W.A., Shao, X., Morin, A., Siroux, V., Kwan, T., Ge, B., Aissi, D., Chen, L., Vasquez, L., Allum, F., et al. (2017). Functional variation in allelic methylomes underscores a strong genetic contribution and reveals novel epigenetic alterations in the human epigenome. Genome Biol 18, 50.

30. Gluckman, P.D., Hanson, M.A., Cooper, C., and Thornburg, K.L. (2008). Effect of in utero and early-life conditions on adult health and disease. N Engl J Med 359, 61–73.

31. Klengel, T., Mehta, D., Anacker, C., Rex-Haffner, M., Pruessner, J.C., Pariante, C.M., Pace, T.W., Mercer, K.B., Mayberg, H.S., Bradley, B., et al. (2013). Allele-specific FKBP5 DNA demethylation mediates gene-childhood trauma interactions. Nat Neurosci 16, 33–41.

32. Chen, L., Pan, H., Tuan, T.A., Teh, A.L., MacIsaac, J.L., Mah, S.M., McEwen, L.M., Li, Y., Chen, H., Broekman, B.F., et al. (2015). Brain-derived neurotrophic factor (BDNF) Val66Met polymorphism influences the association of the methylome with maternal anxiety and neonatal brain volumes. Dev Psychopathol 27, 137–150.

33. Gonseth, S., de Smith, A.J., Roy, R., Zhou, M., Lee, S.T., Shao, X., Ohja, J., Wrensch, M.R., Walsh, K.M., Metayer, C., et al. (2016). Genetic contribution to variation in DNA methylation at maternal smoking-sensitive loci in exposed neonates. Epigenetics 11, 664–673.

34. Teh, A.L., Pan, H., Chen, L., Ong, M.L., Dogra, S., Wong, J., MacIsaac, J.L., Mah, S.M., McEwen, L.M., Saw, S.M., et al. (2014). The effect of genotype and in utero environment on interindividual variation in neonate DNA methylomes. Genome Res 24, 1064–1074.

35. Girchenko, P., Hamalainen, E., Kajantie, E., Pesonen, A.K., Villa, P., Laivuori, H., and Raikkonen, K. (2016). Prediction and Prevention of Preeclampsia and Intrauterine Growth Restriction (PREDO) study. Int J Epidemiol.

36. Graham, A.M., Rasmussen, J.M., Rudolph, M.D., Heim, C.M., Gilmore, J.H., Styner, M., Potkin, S.G., Entringer, S., Wadhwa, P.D., Fair, D.A., et al. (2018). Maternal Systemic Interleukin-6 During Pregnancy Is Associated With Newborn Amygdala Phenotypes and Subsequent Behavior at 2 Years of Age. Biol Psychiatry 83, 109–119.

37. Moog, N.K., Entringer, S., Rasmussen, J.M., Styner, M., Gilmore, J.H., Kathmann, N., Heim, C.M., Wadhwa, P.D., and Buss, C. (2018). Intergenerational Effect of Maternal Exposure to Childhood Maltreatment on Newborn Brain Anatomy. Biol Psychiatry 83, 120–127.

38. Entringer, S., Buss, C., Rasmussen, J.M., Lindsay, K., Gillen, D.L., Cooper, D.M., and Wadhwa, P.D. (2017). Maternal Cortisol During Pregnancy and Infant Adiposity: A Prospective Investigation. J Clin Endocrinol Metab 102, 1366–1374.

39. Stein, D.J., Koen, N., Donald, K.A., Adnams, C.M., Koopowitz, S., Lund, C., Marais, A., Myers, B., Roos, A., Sorsdahl, K., et al. (2015). Investigating the psychosocial determinants of child health in Africa: The Drakenstein Child Health Study. J Neurosci Methods 252, 27–35.

40. Zar, H.J., Barnett, W., Myer, L., Stein, D.J., and Nicol, M.P. (2015). Investigating the early-life determinants of illness in Africa: the Drakenstein Child Health Study. Thorax 70, 592–594.

41. Ronningen, K.S., Paltiel, L., Meltzer, H.M., Nordhagen, R., Lie, K.K., Hovengen, R., Haugen, M., Nystad, W., Magnus, P., and Hoppin, J.A. (2006). The biobank of the Norwegian Mother and Child Cohort Study: a resource for the next 100 years. Eur J Epidemiol 21, 619–625.

42. Zhou, J., and Troyanskaya, O.G. (2015). Predicting effects of noncoding variants with deep learning-based sequence model. Nat Methods 12, 931–934.

43. Radloff, L.S. (1977). The CES-D Scale: A Self-Report Depression Scale for Research in the General Population. Appl Psychol Meas 1, 385–401.

44. Spielberger, C.D. (1989). State-Trait Anxiety Inventory: Bibliography.(Consulting Psychologists Press).

45. Aryee, M.J., Jaffe, A.E., Corrada-Bravo, H., Ladd-Acosta, C., Feinberg, A.P., Hansen, K.D., and Irizarry, R.A. (2014). Minfi: a flexible and comprehensive Bioconductor package for the analysis of Infinium DNA methylation microarrays. Bioinformatics 30, 1363–1369.

46. Morin, A.M., Gatev, E., McEwen, L.M., MacIsaac, J.L., Lin, D.T.S., Koen, N., Czamara, D., Raikkonen, K., Zar, H.J., Koenen, K., et al. (2017). Maternal blood contamination of collected cord blood can be identified using DNA methylation at three CpGs. Clin Epigenetics 9, 75.

47. Fortin, J.P., Labbe, A., Lemire, M., Zanke, B.W., Hudson, T.J., Fertig, E.J., Greenwood, C.M., and Hansen, K.D. (2014). Functional normalization of 450k methylation array data improves replication in large cancer studies. Genome Biol 15, 503.

48. Johnson, W.E., Li, C., and Rabinovic, A. (2007). Adjusting batch effects in microarray expression data using empirical Bayes methods. Biostatistics 8, 118–127.

49. Leek, J.T., Johnson, W.E., Parker, H.S., Jaffe, A.E., and Storey, J.D. (2012). The sva package for removing batch effects and other unwanted variation in high-throughput experiments. Bioinformatics 28, 882–883.

50. Chen, Y.A., Lemire, M., Choufani, S., Butcher, D.T., Grafodatskaya, D., Zanke, B.W., Gallinger, S., Hudson, T.J., and Weksberg, R. (2013). Discovery of cross-reactive probes and polymorphic CpGs in the Illumina Infinium HumanMethylation450 microarray. Epigenetics 8, 203–209.

51. Price, M.E., Cotton, A.M., Lam, L.L., Farre, P., Emberly, E., Brown, C.J., Robinson, W.P., and Kobor, M.S. (2013). Additional annotation enhances potential for biologically-relevant analysis of the Illumina Infinium HumanMethylation450 BeadChip array. Epigenetics Chromatin 6, 4.

52. McCartney, D.L., Walker, R.M., Morris, S.W., McIntosh, A.M., Porteous, D.J., and Evans, K.L. (2016). Identification of polymorphic and off-target probe binding sites on the Illumina Infinium MethylationEPIC BeadChip. Genom Data 9, 22–24.

53. Bakulski, K.M., Feinberg, J.I., Andrews, S.V., Yang, J., Brown, S., S, L.M., Witter, F., Walston, J., Feinberg, A.P., and Fallin, M.D. (2016). DNA methylation of cord blood cell types: Applications for mixed cell birth studies. Epigenetics 11, 354–362.

54. Ong, M.L., and Holbrook, J.D. (2014). Novel region discovery method for Infinium 450K DNA methylation data reveals changes associated with aging in muscle and neuronal pathways. Aging Cell 13, 142–155.

55. van der Westhuizen, C., Wyatt, G., Williams, J.K., Stein, D.J., and Sorsdahl, K. (2016). Validation of the Self Reporting Questionnaire 20-Item (SRQ-20) for Use in a Low- and Middle-Income Country Emergency Centre Setting. Int J Ment Health Addict 14, 37–48.

56. Beck, A.T., Ward, C.H., Mendelson, M., Mock, J., and Erbaugh, J. (1961). An inventory for measuring depression. Arch Gen Psychiatry 4, 561–571.

57. Group, W.A.W. (2002). The Alcohol, Smoking and Substance Involvement Screening Test (ASSIST): development, reliability and feasibility. Addiction 97, 1183–1194.

58. Magnus, P., Birke, C., Vejrup, K., Haugan, A., Alsaker, E., Daltveit, A.K., Handal, M., Haugen, M., Hoiseth, G., Knudsen, G.P., et al. (2016). Cohort Profile Update: The Norwegian Mother and Child Cohort Study (MoBa). Int J Epidemiol 45, 382–388.

59. Haberg, S.E., London, S.J., Nafstad, P., Nilsen, R.M., Ueland, P.M., Vollset, S.E., and Nystad, W. (2011). Maternal folate levels in pregnancy and asthma in children at age 3 years. J Allergy Clin Immunol 127, 262–264, 264 e261.

60. Joubert, B.R., Haberg, S.E., Nilsen, R.M., Wang, X., Vollset, S.E., Murphy, S.K., Huang, Z., Hoyo, C., Midttun, O., Cupul-Uicab, L.A., et al. (2012). 450K epigenome-wide scan identifies differential DNA methylation in newborns related to maternal smoking during pregnancy. Environ Health Perspect 120, 1425–1431.

61. Irgens, L.M. (2000). The Medical Birth Registry of Norway. Epidemiological research and surveillance throughout 30 years. Acta Obstet Gynecol Scand 79, 435–439.

62. Bibikova, M., Barnes, B., Tsan, C., Ho, V., Klotzle, B., Le, J.M., Delano, D., Zhang, L., Schroth, G.P., Gunderson, K.L., et al. (2011). High density DNA methylation array with single CpG site resolution. Genomics 98, 288–295.

63. Teschendorff, A.E., Marabita, F., Lechner, M., Bartlett, T., Tegner, J., Gomez-Cabrero, D., and Beck, S. (2013). A beta-mixture quantile normalization method for correcting probe design bias in Illumina Infinium 450 k DNA methylation data. Bioinformatics 29, 189–196.

64. Akaike, H. (1973). Information theory and an extension of the maximum likelihood principle. In Proceedings of the Second International Symposium on Information Theory (ed (Budapest, Hungary, Akademiai Kiado), pp 267–281.

65. Griffon, A., Barbier, Q., Dalino, J., van Helden, J., Spicuglia, S., and Ballester, B. (2015). Integrative analysis of public ChIP-seq experiments reveals a complex multi-cell regulatory landscape. Nucleic Acids Res 43, e27.

66. Consortium, E.P. (2004). The ENCODE (ENCyclopedia Of DNA Elements) Project. Science 306, 636–640.

67. Gu, J., Stevens, M., Xing, X., Li, D., Zhang, B., Payton, J.E., Oltz, E.M., Jarvis, J.N., Jiang, K., Cicero, T., et al. (2016). Mapping of Variable DNA Methylation Across Multiple Cell Types Defines a Dynamic Regulatory Landscape of the Human Genome. G3 (Bethesda) 6, 973–986.

68. Bonder, M.J., Luijk, R., Zhernakova, D.V., Moed, M., Deelen, P., Vermaat, M., van Iterson, M., van Dijk, F., van Galen, M., Bot, J., et al. (2017). Disease variants alter transcription factor levels and methylation of their binding sites. Nat Genet 49, 131–138.

69. Smith, A.K., Kilaru, V., Kocak, M., Almli, L.M., Mercer, K.B., Ressler, K.J., Tylavsky, F.A., and Conneely, K.N. (2014). Methylation quantitative trait loci (meQTLs) are consistently detected across ancestry, developmental stage, and tissue type. BMC Genomics 15, 145.

70. Wagner, J.R., Busche, S., Ge, B., Kwan, T., Pastinen, T., and Blanchette, M. (2014). The relationship between DNA methylation, genetic and expression inter-individual variation in untransformed human fibroblasts. Genome Biol 15, R37.

71. Autism Spectrum Disorders Working Group of The Psychiatric Genomics, C. (2017). Meta-analysis of GWAS of over 16,000 individuals with autism spectrum disorder highlights a novel locus at 10q24.32 and a significant overlap with schizophrenia. Mol Autism 8, 21.

72. Demontis, D., Walters, R., Martina, J., Mattheisen, M., Als, T., Agerbo, E., and Belliveau, R. (2017). Discovery of the first genome-wide significant risk loci for ADHD. BioRxiv.

73. Psychiatric, G.C.B.D.W.G. (2011). Large-scale genome-wide association analysis of bipolar disorder identifies a new susceptibility locus near ODZ4. Nat Genet 43, 977–983.

74. Wray, N.R., Ripke, S., Mattheisen, M., Trzaskowski, M., Byrne, E.M., Abdellaoui, A., Adams, M.J., Agerbo, E., Air, T.M., Andlauer, T.M.F., et al. (2018). Genome-wide association analyses identify 44 risk variants and refine the genetic architecture of major depression. Nat Genet 50, 668–681.

75. Schizophrenia Working Group of the Psychiatric Genomics, C. (2014). Biological insights from 108 schizophrenia-associated genetic loci. Nature 511, 421–427.

76. Cross-Disorder Group of the Psychiatric Genomics, C. (2013). Identification of risk loci with shared effects on five major psychiatric disorders: a genome-wide analysis. Lancet 381, 1371–1379.

77. Liu, J.Z., van Sommeren, S., Huang, H., Ng, S.C., Alberts, R., Takahashi, A., Ripke, S., Lee, J.C., Jostins, L., Shah, T., et al. (2015). Association analyses identify 38 susceptibility loci for inflammatory bowel disease and highlight shared genetic risk across populations. Nat Genet 47, 979–986.

78. Morris, A.P., Voight, B.F., Teslovich, T.M., Ferreira, T., Segre, A.V., Steinthorsdottir, V., Strawbridge, R.J., Khan, H., Grallert, H., Mahajan, A., et al. (2012). Large-scale association analysis provides insights into the genetic architecture and pathophysiology of type 2 diabetes. Nat Genet 44, 981–990.

79. Horikoshi, M., Mgi, R., van de Bunt, M., Surakka, I., Sarin, A.P., Mahajan, A., Marullo, L., Thorleifsson, G., Hgg, S., Hottenga, J.J., et al. (2015). Discovery and Fine-Mapping of Glycaemic and Obesity-Related Trait Loci Using High-Density Imputation. PLoS Genet 11, e1005230.

80. Crudo, A., Petropoulos, S., Suderman, M., Moisiadis, V.G., Kostaki, A., Hallett, M., Szyf, M., and Matthews, S.G. (2013). Effects of antenatal synthetic glucocorticoid on glucocorticoid receptor binding, DNA methylation, and genome-wide mRNA levels in the fetal male hippocampus. Endocrinology 154, 4170–4181.

81. Glad, C.A., Andersson-Assarsson, J.C., Berglund, P., Bergthorsdottir, R., Ragnarsson, O., and Johannsson, G. (2017). Reduced DNA methylation and psychopathology following endogenous hypercortisolism - a genome-wide study. Sci Rep 7, 44445.

82. Kress, C., Thomassin, H., and Grange, T. (2006). Active cytosine demethylation triggered by a nuclear receptor involves DNA strand breaks. Proc Natl Acad Sci U S A 103, 11112–11117.

83. Seifuddin, F., Wand, G., Cox, O., Pirooznia, M., Moody, L., Yang, X., Tai, J., Boersma, G., Tamashiro, K., Zandi, P., et al. (2017). Genome-wide Methyl-Seq analysis of blood-brain targets of glucocorticoid exposure. Epigenetics 12, 637–652.

84. Markunas, C.A., Wilcox, A.J., Xu, Z., Joubert, B.R., Harlid, S., Panduri, V., Haberg, S.E., Nystad, W., London, S.J., Sandler, D.P., et al. (2016). Maternal Age at Delivery Is Associated with an Epigenetic Signature in Both Newborns and Adults. PLoS One 11, e0156361.

85. Moran, S., Arribas, C., and Esteller, M. (2016). Validation of a DNA methylation microarray for 850,000 CpG sites of the human genome enriched in enhancer sequences. Epigenomics 8, 389–399.

86. Pidsley, R., Zotenko, E., Peters, T.J., Lawrence, M.G., Risbridger, G.P., Molloy, P., Van Djik, S., Muhlhausler, B., Stirzaker, C., and Clark, S.J. (2016). Critical evaluation of the Illumina MethylationEPIC BeadChip microarray for whole-genome DNA methylation profiling. Genome Biol 17, 208.

87. Sandoval, J., Heyn, H., Moran, S., Serra-Musach, J., Pujana, M.A., Bibikova, M., and Esteller, M. (2011). Validation of a DNA methylation microarray for 450,000 CpG sites in the human genome. Epigenetics 6, 692–702.

88. Mehta, D., Klengel, T., Conneely, K.N., Smith, A.K., Altmann, A., Pace, T.W., Rex-Haffner, M., Loeschner, A., Gonik, M., Mercer, K.B., et al. (2013). Childhood maltreatment is associated with distinct genomic and epigenetic profiles in posttraumatic stress disorder. Proc Natl Acad Sci U S A 110, 8302–8307.

89. Grishkevich, V., Yanai, I. (2013). The genomic determinants of genotype × environment interactions in gene expression. Trends in Genetics 29, 479–487.

90. Grishkevich, V., Ben-Elazar, S., Hashimshony, T., Schott, D.H., Hunter, C.P., and Yanai, I. (2012). A genomic bias for genotype-environment interactions in C. elegans. Mol Syst Biol 8, 587.

91. Feinberg, A.P., and Irizarry, R.A. (2010). Evolution in health and medicine Sackler colloquium: Stochastic epigenetic variation as a driving force of development, evolutionary adaptation, and disease. Proc Natl Acad Sci U S A 107 Suppl 1, 1757–1764.

92. Elliott, G., Hong, C., Xing, X., Zhou, X., Li, D., Coarfa, C., Bell, R.J., Maire, C.L., Ligon, K.L., Sigaroudinia, M., et al. (2015). Intermediate DNA methylation is a conserved signature of genome regulation. Nat Commun 6, 6363.

93. Zhang, P. (1992). Inference after variable selection in linear regression models. Biometrika 79, 741–746.

